# Ultrapotent SARS-CoV-2 neutralizing antibodies with protective efficacy against newly emerged mutational variants

**DOI:** 10.1101/2021.04.19.440481

**Authors:** Tingting Li, Xiaojian Han, Chenjian Gu, Hangtian Guo, Huajun Zhang, Yingming Wang, Chao Hu, Kai Wang, Fengjiang Liu, Feiyang Luo, Yanan Zhang, Jie Hu, Wang Wang, Shenglong Li, Yanan Hao, Meiying Shen, Jingjing Huang, Yingyi Long, Shuyi Song, Ruixin Wu, Song Mu, Qian Chen, Fengxia Gao, Jianwei Wang, Shunhua Long, Luo Li, Yang Wu, Yan Gao, Wei Xu, Xia Cai, Di Qu, Zherui Zhang, Hongqing Zhang, Na Li, Qingzhu Gao, Guiji Zhang, Changlong He, Wei Wang, Xiaoyun Ji, Ni Tang, Zhenghong Yuan, Youhua Xie, Haitao Yang, Bo Zhang, Ailong Huang, Aishun Jin

**Affiliations:** Department of Immunology, College of Basic Medicine, Chongqing Medical University, Chongqing, 400010, China; Chongqing Key Laboratory of Basic and Translational Research of Tumor Immunology, Chongqing Medical University, Chongqing, 400010, China; Key Laboratory of Medical Molecular Virology, Department of Medical Microbiology and Parasitology, School of Basic Medical Sciences, Shanghai Medical College, Fudan University, Shanghai, 200032, China; Shanghai Institute for Advanced Immunochemical Studies and School of Life Science and Technology, ShanghaiTech University, Shanghai, 201210, China; State Key Laboratory of Pharmaceutical Biotechnology, School of Life Sciences, Nanjing University, Nanjing, Jiangsu, 210023, China; State Key Laboratory of Virology, Wuhan Institute of Virology, Center for Biosafety Mega-Science, Chinese Academy of Sciences, Wuhan, 430071, China; Key Laboratory of Molecular Biology on Infectious Diseases, Ministry of Education, Chongqing Medical University, Chongqing, 400010, China; Key Laboratory of Special Pathogens and Biosafety, Wuhan Institute of Virology, Center for Biosafety Mega-Science, Chinese Academy of Sciences, Wuhan, 430071, China; University of Chinese Academy of Sciences, Beijing, 100049, China; Department of Breast Surgery, Harbin Medical University Cancer Hospital, Harbin, 150000, China; Institute of life sciences, Chongqing Medical University, Chongqing, 400010, China

**Author notes:** These authors contributed equally (Tingting Li, Xiaojian Han, Chenjian Gu, Hangtian Guo, Huajun Zhang, Yingming Wang). Correspondence (Y.H.X); (H.T.Y); (B.Z); (A.L.H); (A.S.J).

## Abstract

Accumulating mutations in the SARS-CoV-2 Spike (S) protein can increase the possibility of immune escape, challenging the present COVID-19 prophylaxis and clinical interventions. Here, 3 receptor binding domain (RBD) specific monoclonal antibodies (mAbs), 58G6, 510A5 and 13G9, with high neutralizing potency blocking authentic SARS-CoV-2 virus displayed remarkable efficacy against authentic B.1.351 virus. Each of these 3 mAbs in combination with one neutralizing Ab recognizing non-competing epitope exhibited synergistic effect against authentic SARS-CoV-2 virus. Surprisingly, structural analysis revealed that 58G6 and 13G9, encoded by the *IGHV1-58* and the *IGKV3-20* germline genes, both recognized the steric region S^470-495^ on the RBD, overlapping the E484K mutation presented in B.1.351. Also, 58G6 directly bound to another region S^450-458^ in the RBD. Significantly, 58G6 and 510A5 both demonstrated prophylactic efficacy against authentic SARS-CoV-2 and B.1.351 viruses in the transgenic mice expressing human ACE2 (hACE2), protecting weight loss and reducing virus loads. These 2 ultrapotent neutralizing Abs can be promising candidates to fulfill the urgent needs for the prolonged COVID-19 pandemic.

## Introduction

The persistence of COVID-19 in the global population can result in the accumulation of specific mutations of SARS-CoV-2 with increased infectivity and/or reduced susceptibility to neutralization^1–12^. Highly transmissible SARS-CoV-2 variants, such as B.1.351 emerged in South Africa, harbor multiple immune escape mutations, and have raised global concerns for the efficacy of available interventions and for re-infection^2–9,11^. As these challenges presented, the protective efficacy of current antibody-based countermeasures needs to be thoroughly assessed against the current mutational variants.

The major interest of neutralizing therapies has been targeted towards SARS-CoV-2 RBD, which is the core region for the host cell receptor ACE2 engagement^13–23^. B.1.351 bears 3 mutations, S^K417N^, S^E484K^ and S^N501Y^, in its RBD, the first 2 of which have been proven to be the cause for its evasion from neutralizing Ab and serum responses^2–9^. Nevertheless, a small group of SARS-CoV-2 RBD specific neutralizing Abs demonstrated undisturbed *in vitro* potency against B.1.351^2,4–7,9^. Evaluating their therapeutic efficacy against the circulating strains is necessary for the reformulation of protective interventions and vaccines against the evolving pandemic.

Here, we focused on 20 neutralizing Abs selected from a SARS-CoV-2 RBD specific mAb reservoir and confirmed their potency against authentic SARS-CoV-2 virus. Excitingly, at least 3 of our mAbs showed remarkable neutralizing efficacy against authentic B.1.351 virus. 58G6, one of our top neutralizing Abs, was found to target a region of S^450-458^ and a steric site S^470-495^ on the receptor binding motif (RBM). Furthermore, ultrapotent 58G6 and 510A5 exhibited strong prophylactic efficacy in SARS-CoV-2- and B.1.351-infected hACE2-transgenic mice. Our study has characterized a pair of neutralizing Abs with potential effective therapeutic value in clinical applications, which may provide updated information for RBD specific mAbs against the prolonged COVID-19 pandemic.

## Results

### SARS-CoV-2 RBD specific neutralizing Abs exhibited sustained efficacy against authentic B.1.351

By our recently established rapid neutralizing Abs screening system^24^, we have successfully obtained 20 neutralizing Abs with high affinities to SARS-CoV-2 RBD from COVID-19 convalescent individuals, and their neutralizing potencies were confirmed by the half inhibition concentrations (IC_50_s) against authentic SARS-CoV-2 virus quantified via qRT-PCR (Fig. 1a, c and Extended Data Fig. 1). Here, we analyzed the neutralizing potency of our top 10 neutralizing Abs against authentic SARS-CoV-2 and B.1.351 viruses by the plaque-reduction neutralization testing (PRNT). At least 3 of our potent neutralizing Abs 58G6, 510A5 and 13G9 exhibited striking neutralizing efficacy against SARS-CoV-2, with the IC_50_s value ranging from 1.285 to 9.174 ng/mL (Fig. 1b, c). Importantly, the RBD escape mutations of B.1.351 did not compromise the neutralizing efficacy of 58G6 and 510A5, with the IC_50_s of 1.660 and 2.235 ng/ml respectively (Fig. 1b, c). As reported for a wide range of RBD specific neutralizing Abs^2–9^, authentic B.1.351 virus has challenged some of the tested mAbs (Fig. 1b, c). However, majority of our top 10 mAbs still exhibited neutralizing capabilities against this variant (Fig. 1b, c). Of note, the neutralizing potencies of all 10 mAbs against the B. 1.1.7 pseudovirus were shown to be similar to those against the SARS-CoV-2 pseudovirus (Fig. 1c and Extended Data Fig. 2). In addition, the binding affinity of 58G6 to the B.1.351 S1 subunit was comparable to that to the SARS-CoV-2 S1, while 510A5 and 13G9 showed higher binding affinity to the S1 subunit of SARS-CoV-2 than that of B.1.351 (Extended Data Fig. 3). Majority of these top 20 neutralizing Abs exhibited no cross-reactivity to the SARS-CoV S protein or the MARS-CoV S protein (Extended Data Fig. 4). Collectively, 3 RBD specific mAbs demonstrated potent neutralizing efficacy against authentic SARS-CoV-2 and B.1.351 viruses, suggesting that our neutralizing Abs might be applied for the current COVID-19 pandemic.

**Fig. 1.**
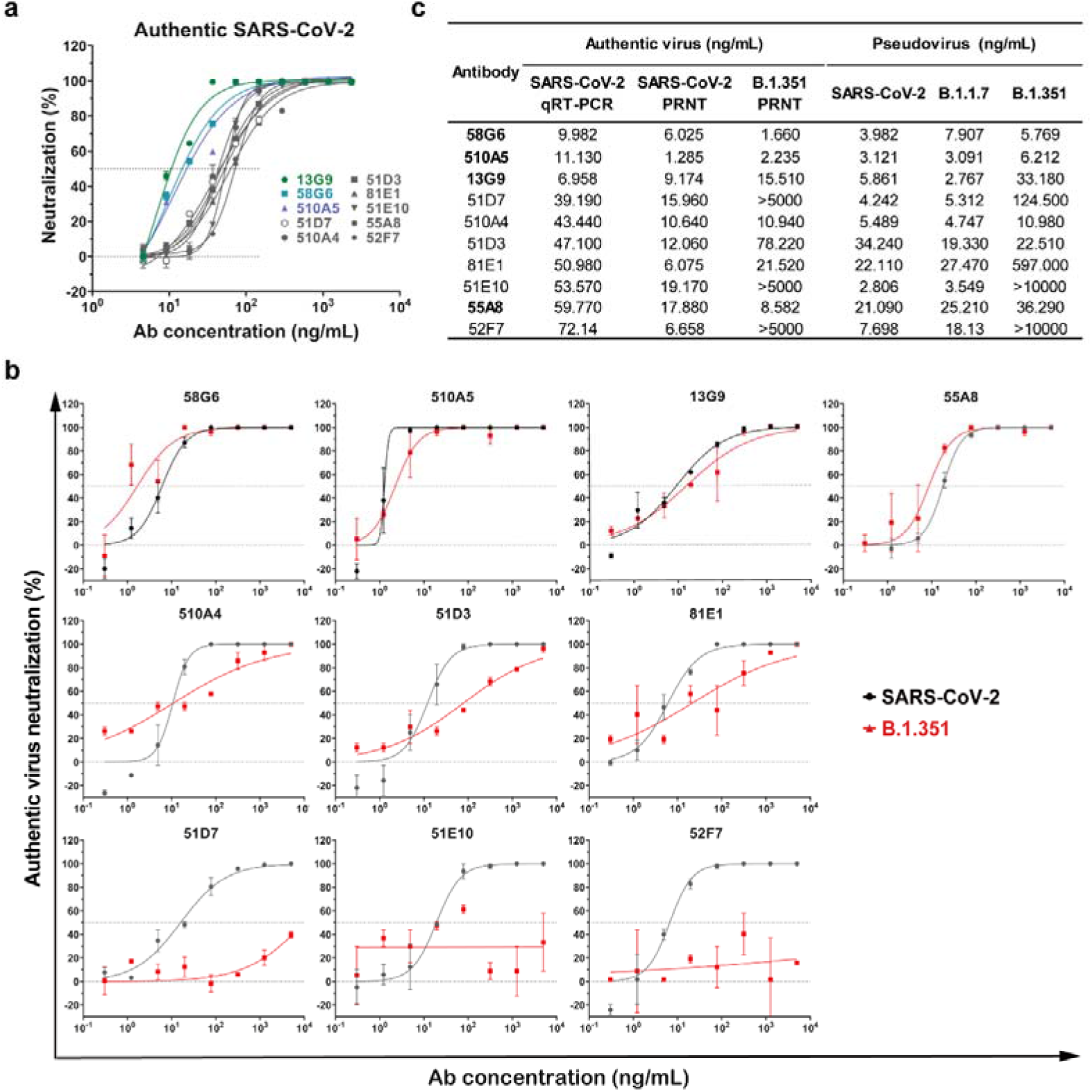
The neutralizing capabilities of the top 10 mAbs against authentic SARS-CoV-2 and B.1.351 viruses. The neutralizing potency of the top 10 mAbs was measured by authentic SARS-CoV-2 (nCoV-SH01) neutralization assay and quantified by qRT-PCR (a) or authentic SARS-CoV-2 (WIV04) and B.1.351 neutralization assays and quantified by PRNT (b). The IC_50_s were summarized in (c). Dashed line indicated 0% or 50% reduction in viral neutralization. Data for each mAb were obtained from a representative neutralization experiment, with at least two replicates, presented as mean ± SEM. Effective Abs against authentic B.1.351 were shown in bold.

### The epitopes for potent neutralizing Abs overlapped a key site on SARS-CoV-2 RBD

To define potential antigenic sites on SARS-CoV-2 RBD, we performed competitive ELISA with the above top 20 neutralizing Abs and the other 54 mAbs selected from our developed RBD-specific mAb reservoir. As shown in Fig. 2a, 5 groups of mAbs were identified according to their recognition sites, each of which consisted of mAbs competing for the epitope for 13G9 (13G9e), the epitope recognized by a non-neutralizing SARS-CoV-2 specific mAb 81A11 (81A11e), or the epitope recognized by a SARS-CoV specific neutralizing Ab CR3022 (CR3022e) (Fig. 2a). Interestingly, the epitopes recognized by the majority of potent neutralizing Abs overlapped with 13G9e (Fig. 2a). Next, we confirmed that the top 20 mAbs could directly inhibit the interaction of SARS-CoV-2 RBD and ACE2 by the competitive ELISA and surface plasmon resonance (SPR) assay (Extended Data Fig. 5 and 6). To assess the interrelationships between the epitopes recognized by our top 20 neutralizing Abs in detail, we performed competitive ELISA using biotinylated mAbs. We found that 16 of them competed with 13G9, whereas the antigenic sites of the other 4 Abs (510A5, 55A8, 57F7 and 07C1) overlapped with an independent epitope (510A5e) (Extended Data Fig. 7). These findings suggest that there are at least 2 independent epitopes on the RBD related to SARS-CoV-2 neutralization, from which 13G9e may represent a key antigenic site for the binding of potent neutralizing Abs to the RBD.

**Fig. 2.**
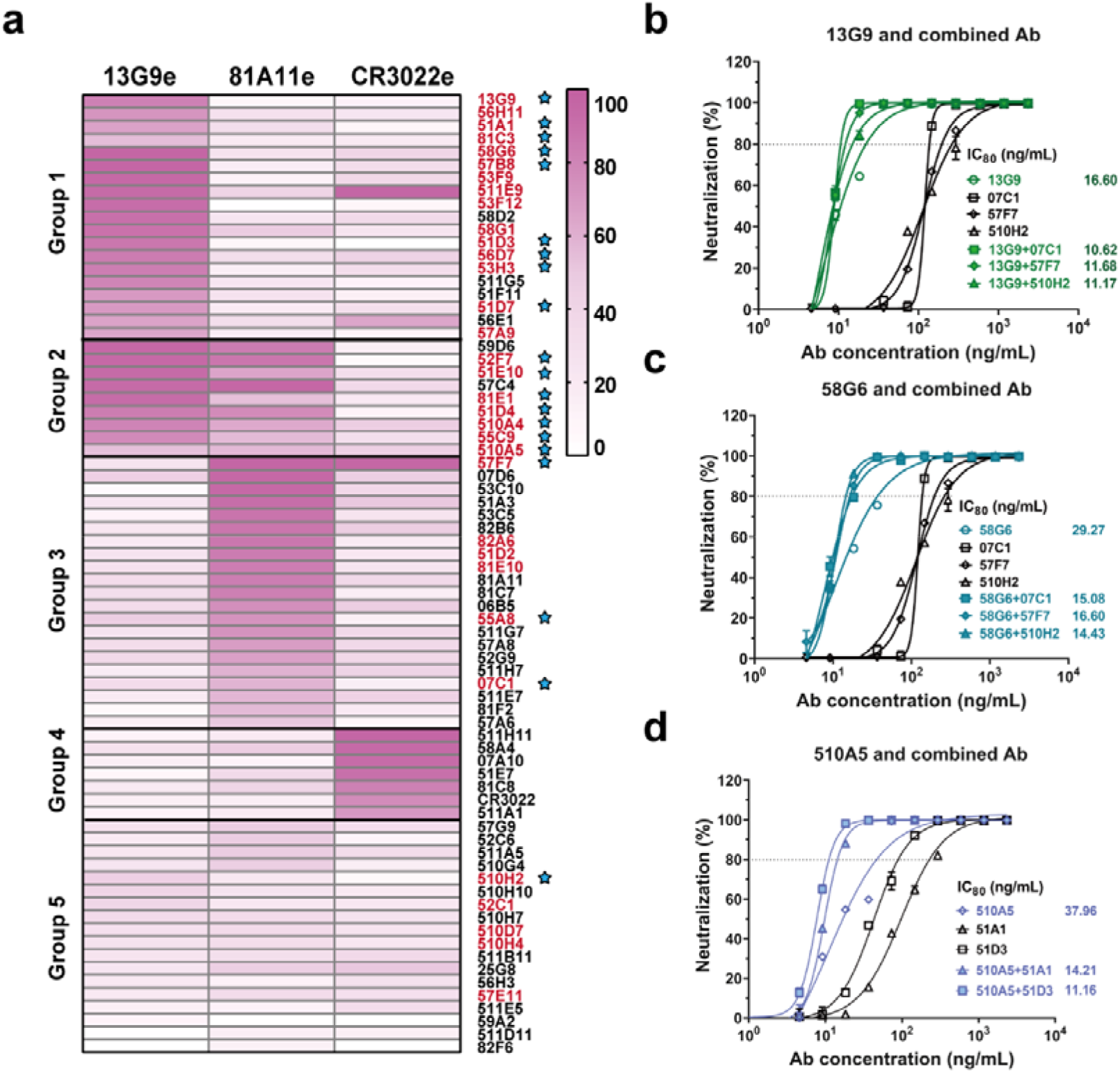
Epitope mapping of mAbs and the analysis of neutralizing Abs from different groups. (a) Epitope mapping of purified mAbs targeting three independent epitopes (13G9e, 81A11e and CR3022e). All mAbs in Group 1 competed with 13G9; each mAb in Group 3 competed with 81A11; Group 2 consisted of mAbs cross-reacted with 13G9e and 81A11e, the latter to a lesser extent; all mAbs in Group 4 targeted the epitopes overlapping with CR3022e, and the mAbs in Group 5 recognized none of these 3 epitopes. All neutralizing Abs identified by authentic SARS-CoV-2 CPE assay were labelled in red. The top 20 mAbs identified by qRT-PCR with authentic SARS-CoV-2 virus were indicated by blue stars. The synergistic effects of 13G9 (b) and 58G6 (c) with 07C1 or 57F7 recognizing 81A11e, or 510H2 with no clearly identified epitope, against authentic SARS-CoV-2 virus were quantified by qRT-PCR. (d) The synergistic effects of 510A5 with 51A1 or 51D3 recognizing 13G9e, against authentic SARS-CoV-2 virus were quantified by qRT-PCR. Dashed line indicated 80% inhibition in the viral infectivity. Data for each mAb were obtained from a representative neutralization experiment of three replicates, presented as mean ± SEM.

To test whether our mAbs could elicit synergistic effect against SARS-CoV-2, we paired each of the top 3 neutralizing Abs (58G6, 510A5 or 13G9) with one Ab exhibiting much lower potency from another group shown in Fig 2a. Synergistic effects were observed for all combinations at higher levels of inhibition against the authentic virus, confirming the synergistic advantage of neutralizing Ab cocktails (Fig. 2b-d). Of note, adding neutralizing Ab from a different cluster barely reduced the IC_50_s of the top 3 mAbs, indicating that our potent mAbs alone were sufficient in neutralizing SARS-CoV-2 (Fig. 2b-d).

### 58G6 recognized a linear binding region in the denatured RBD

To determine the precise interactive regions of our potent neutralizing Abs, first, we assessed the binding ability of the top 20 mAbs to the denatured RBD. In a preliminary screening, 9 mAbs from our top 20 mAbs to SARS-CoV-2 RBD were found to be capable of directly binding to the denatured RBD (Extended Data Fig. 8). Therefore, we designed and synthesized fifteen 20-mer peptides (RBD1 to RBD15), overlapping with 5 amino acids, to cover the entire sequence of the RBD, as amino acids 319-541 of SARS-CoV-2 S (S^319-541^) (Extended Data Fig. 9a). Unexpectedly, instead of a continuous linear region, we found that 5 of these 9 mAbs could simultaneously recognize 3 independent fragments (RBD2, RBD9 and RBD13), while 58G6 only strongly bound to RBD9 (S^439-459^) (Extended Data Fig. 9b, c). To determine the essential amino acid residues in the RBD accounted for 58G6 binding, we re-synthesized two 20-mer peptides overlapping with 15 amino acids (RBD9-1 and RBD9-2), covering the RBD9 specific residues (Extended Data Fig. 9a). The results of peptide ELISA revealed that 58G6 preferentially interacted with RBD9-1 than RBD9, in a dose-dependent manner, whereas no interaction of 58G6 with RBD9-2 was observed (Fig. 3a, b). When we individually replaced each amino acid residue in RBD9-1 (S^444-463^) with alanine (A), we found that the binding of 58G6 to a fragment of 8 amino acids (S^450-457^) was significantly reduced (Fig. 3a). To a lesser extent, S^445-449^ and S^458-463^ also slightly affected the binding of 58G6, and the former might explain for the abolished interaction of 58G6 with RBD9-2 (Fig. 3a). Moreover, we found that RBD9-1 bound to ACE2 in a dose-dependent manner, which could be competitively inhibited by 58G6 (Fig. 3c-e). And the region of S^445-463^ was identified to be critical for the RBD9-1-ACE2 interaction (Fig. 3c, d). Hence, S^445-463^ represents an important region of SARS-CoV-2 RBD for the recognition of neutralizing Abs represented by 58G6. It is worth mentioning that the interaction of 510A5 or 13G9 with the denatured RBD was not observed (Extended Data Fig. 8). Taken together, we evidenced a linear region in the denatured RBD (S^450-457^) that could be recognized by 58G6, which was one of the ultrapotent neutralizing Abs against authentic SARS-CoV-2 and B.1.351 viruses.

**Fig. 3.**
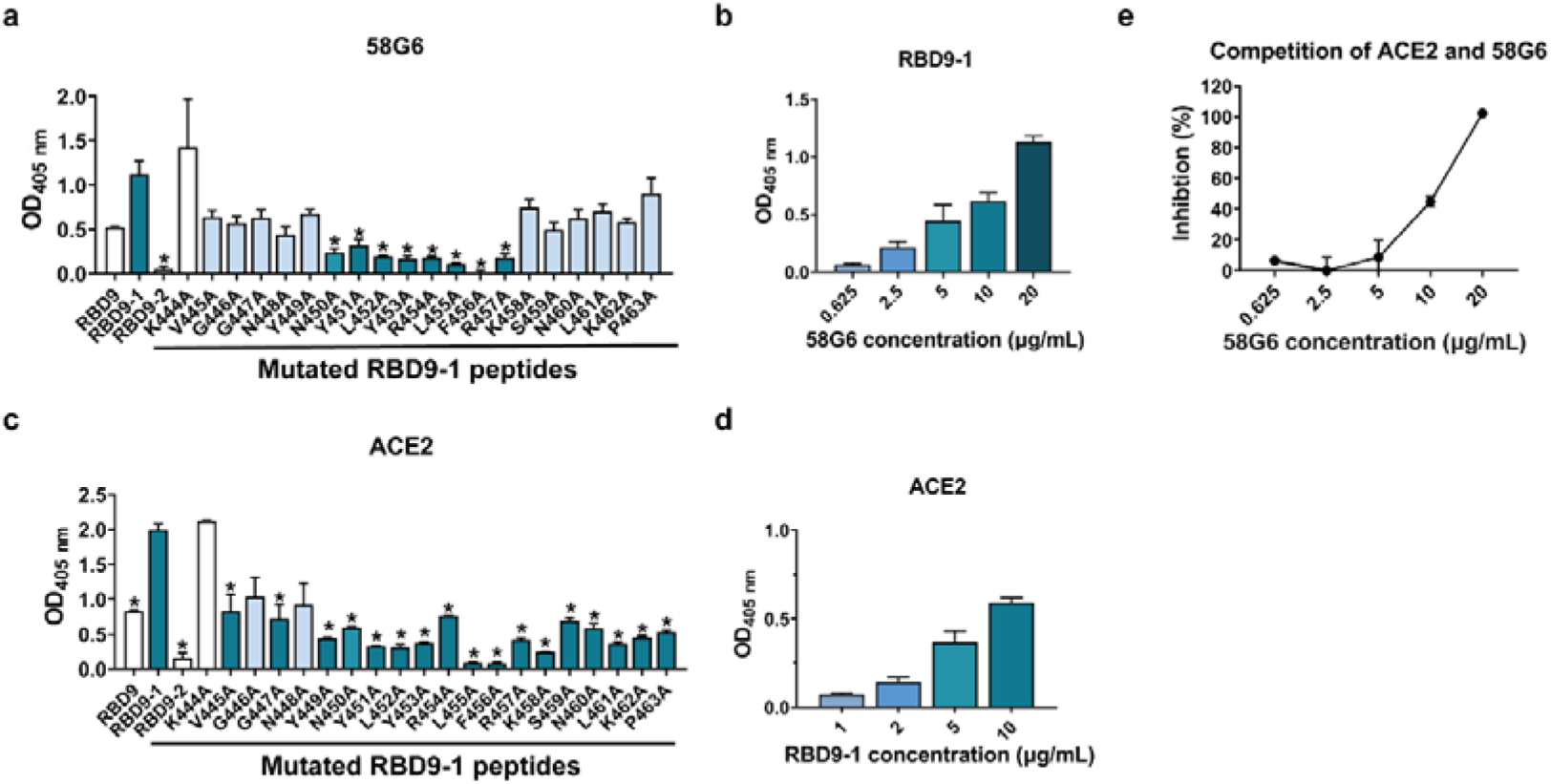
The interaction of 58G6 with a linear region in the denatured RBD. ELISA results of the binding activities of 58G6 (a) or ACE2 (c) to 3 peptides covering sequences in close proximity, RBD9, RBD9-1 and RBD9-2, and single mutations derived from the full length RDB9-1. The binding activity of 58G6 (b) or ACE2 (d) in various concentrations to the RBD9-1 peptide, tested by ELISA. (e) The ability of 58G6 in blocking the interaction between RBD9-1 and ACE2, tested by competitive ELISA. Data are representative of at least 2 independent experiments performed in technical duplicate. The mean ± SEM of duplicates are shown. *, p < 0.05.

### 58G6 and 13G9 encoded by the *IGHV1-58* and *IGKV3-20* germline genes both recognized the steric region of S^470-495^ on the RBD

To further investigate the molecular mechanism of our neutralizing Abs against SARS-CoV-2, we determined the single-particle cryo-electron microscopy (cryo-EM) structures of the antigen binding fragments (Fabs) of 58G6 or 13G9 in complex with the modified SARS-CoV-2 S trimer with stabilizing mutations^25^ (Extended Data Fig. 10a, b). We refined these two complex structures to the overall resolution of 3.6 Å for 58G6 and 3.9 Å for 13G9, respectively (Fig. 4a, b, Extended Data Fig. 10c-j and Extended Data Table. 1). For either the 58G6 or the 13G9 complex, the three-dimensional classification of the cryo-EM data showed the presence of a dominant conformational state of S trimers in complex with the Fabs, with the majority of selected particle images representing a 3-Fab-per-trimer complex (Fig. 4a, b). As shown in Fig. 4a, in individual complex, each 58G6 Fab interacted with one RBD in the “up” state. Similar to the structure of the 58G6 Fab-S complex, only one dominant particle class was observed for the 13G9 Fab-S complex, corresponding to a 3-Fab-bound complex with all 3 RBDs in the “up” conformation (Fig. 4b).

**Fig. 4.**
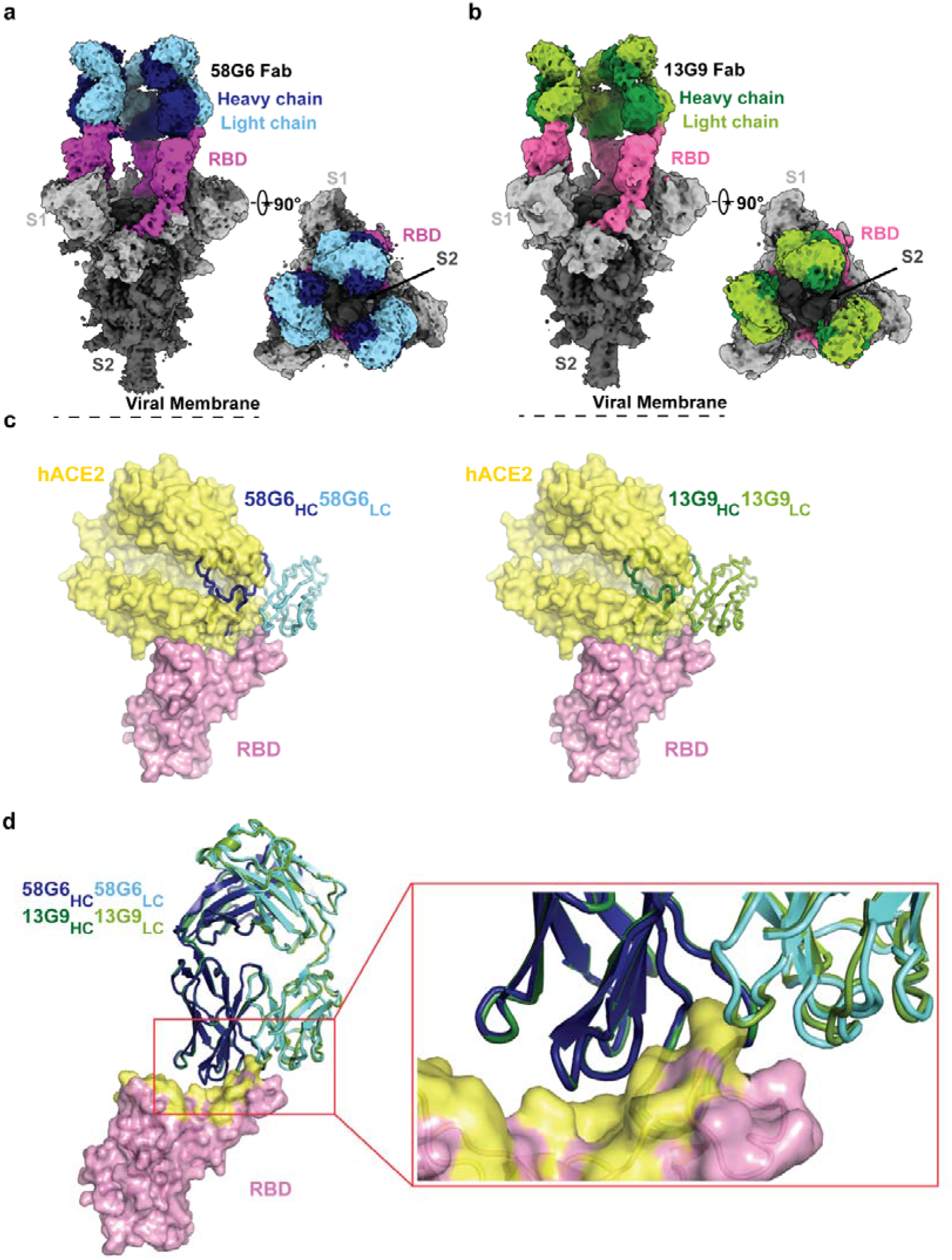
Cryo-EM structures of 58G6 and 13G9 Fabs binding to open S trimer. (a, b) Cryo-EM densities for 58G6 Fab-S (a; 3.6 Å) and 13G9 Fab-S (b; 3.9 Å) complexes, revealing binding of 58G6 or 13G9 to RBDs in the all ‘up’ state. (c) Superimposition of RBD-hACE2 [Protein Data Bank (PDB) ID 6LZG] complex structure together with RBD-58G6 Fab (left) or RBD-13G9 Fab (right) variable domains, respectively. (d) Alignment of 58G6 and 13G9 Fabs on the same RBD. HC, heavy chain; LC, light chain.

Further refinement of the variable domains of 58G6 or 13G9 and the RBD to 3.5 Å or 3.8 Å, respectively, revealed detailed molecular interactions within their binding interface (Extended Data Fig. 10c-f, g-j). These two refined density maps along with the predicted structures of the 58G6 and 13G9 Fabs were used to build the models to illustrate detailed amino acid structures in three dimensions (Extended Data Fig. 11)^26^. Superimposition of the RBDs in the structures of 58G6 Fab-RBD and ACE2-RBD complexes indicates a steric clash between ACE2 and the variable domains on the heavy chain (HC) and the light chain (LC) of 58G6 Fab (Fig. 4c). Such observations indicate that 58G6 can competitively inhibit the interaction between the SARS-CoV-2 RBD and ACE2. Likewise, an almost identical steric clash between 13G9 Fab and ACE2 was observed, indicating that the SARS-CoV-2 RBD-ACE2 interaction can be prohibited by 13G9 (Fig. 4c). When we compared the details of binding interface of these 2 mAbs and RBD, they showed high level of structural similarity (Fig. 4d).

Specifically, majority of the complementarity determining regions (CDRs; CDRH2, CDRH3, CDRL1 and CDRL3) of 58G6 Fab directly participate in the interaction with the steric region of S^470-495^ (Fig. 5a). Meanwhile, 13G9 Fab was shown to recognize the same steric region using its CDRs: CDRH2, CDRH3, CDRL1 and CDRL3 (Fig. 5b). In parallel, an additional site of residues 450-458 on SARS-CoV-2 S (S^450-458^) was observed for 58G6 recognition (Fig. 5b), which contained the linear region of S^450-457^ we had identified with the denatured RBD, as shown above (Fig. 3a, b).

**Fig. 5.**
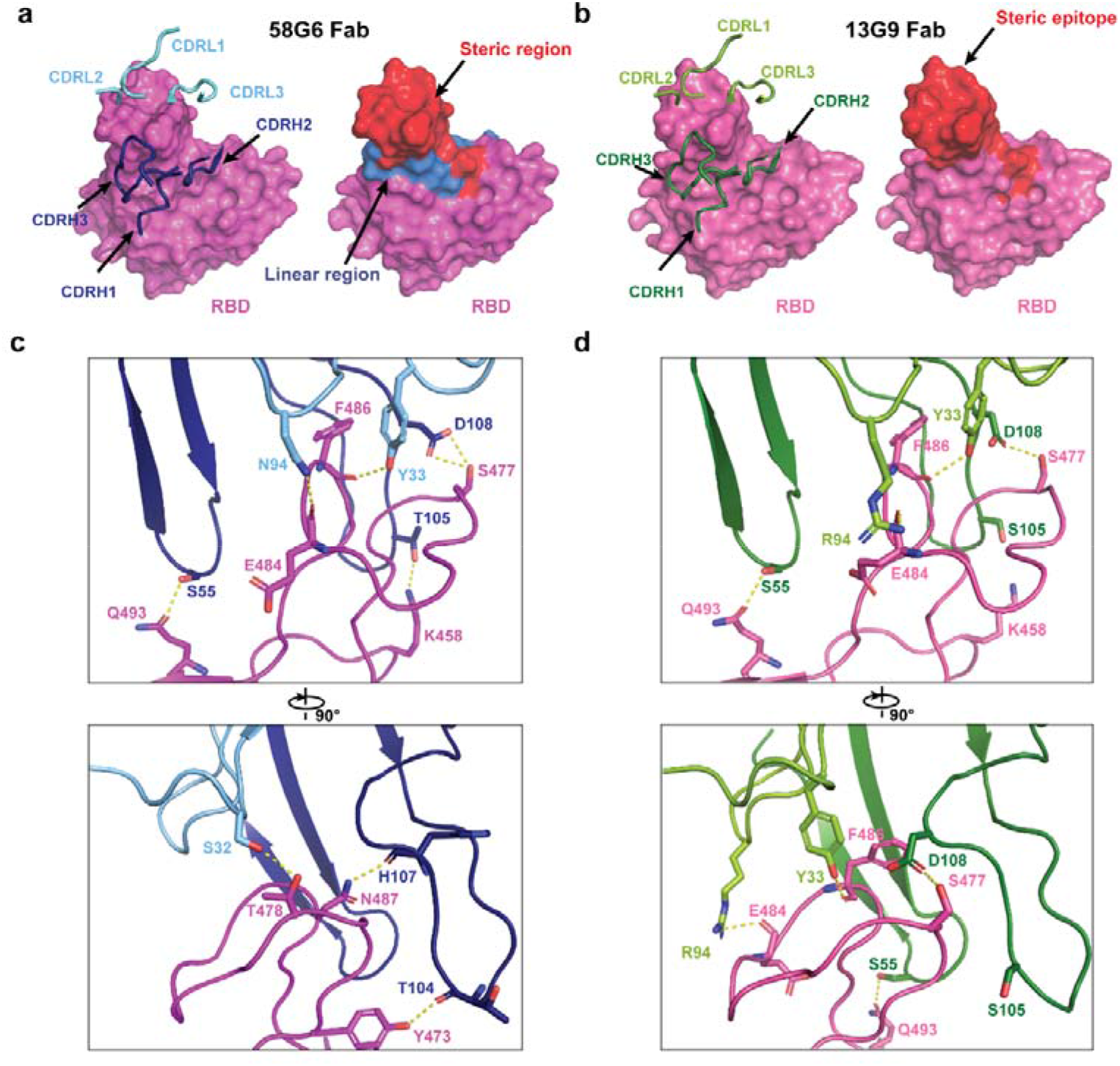
Details of interactions between SARS-CoV-2 RBD and mAbs. (a, b) CDR loops of 58G6 Fab (a, left) and 13G9 Fab (b, left) overlaid on the surface representation of RBD (shown as pink and magenta, respectively), and surface representations of 58G6 epitope (a, right, red and blue) and 13G9 epitope (b, right, red) on the RBD surface. (c, d) The hydrogen bonds at the binding interface between 58G6 (left) or 13G9 (right) and SARS-CoV-2 RBD.

We found that both 58G6 and 13G9 were derived from *IGHV1-58* for the heavy chain and *IGKV3-20* for the light chain, with a few differences in amino acid constitution of their CDRH1, CDRH3 and CDRL3 (Extended Data Table. 2). These identical germline gene origins correlated with the structural similarity between 58G6 and 13G9 (Fig. 4d). Several potential hydrogen bonds were identified on the contact surface of each mAb and RBD, representing the unique network associated with individual CDRs and amino acid residues within the epitope corresponding to each mAb (Fig. 5c, d). In summary, these Fab-S complex structures suggest that 58G6 and 13G9 adopt the same potential neutralizing mechanism, wherein they are capable to simultaneously bind to 3 RBDs, occluding the access of SARS-CoV-2 S to ACE2. Notably, N94 in the CDRL3 of 58G6 or R94 in 13G9 forms a hydrogen bond with the carbonyl group on the main chain, rather than the side chain, of S^E484^ (Fig. 5c, d). Moreover, direct contact with a hydrogen bond was found between T105 in the CDRH3 of 58G6 and K458 in the RBD, but not for S105 in 13G9 (Fig. 5c, d).

### 58G6 and 510A5 showed protective efficacy against SARS-CoV-2 and B.1.351 *in vivo*

Given the IC_50_s of our mAbs 58G6 and 510A5 against authentic B.1.351 were as low as approximately 2 ng/mL *in vitro*, we tested their prophylactic efficacy in the transgenic animal model. Different groups of hACE2 mice received intraperitoneal administration of these 2 mAbs or PBS 24 hours before an intranasal challenge with authentic SARS-CoV-2 (WIV04) or B.1.351. For the hACE2 mice challenged with SARS-CoV-2 (WIV04), the PBS group showed significant loss of body weight, while those animals from either mAb-treated group retained their body weight for 3 days post-infection (Fig. 6a). When challenged with B.1.351, the hACE2 mice receiving PBS showed gradual weight loss and reached an approximately 30% drop at day 3, whereas the treatment of 58G6 or 510A5 effectively stopped the B.1.351-induced weight reduction (Fig. 6a). Importantly, we found that the viral load of either SARS-CoV-2 or B.1.351 in the lung tissues was significantly decreased with a single dose of either mAb (Fig. 6b). These results indicate that 58G6 and 510A5 can effectively protected hACE2 transgenic mice from infectious SARS-CoV-2 and B.1.351, highlighting their prophylactic potential in the present COVID-19 epidemic.

**Fig. 6.**
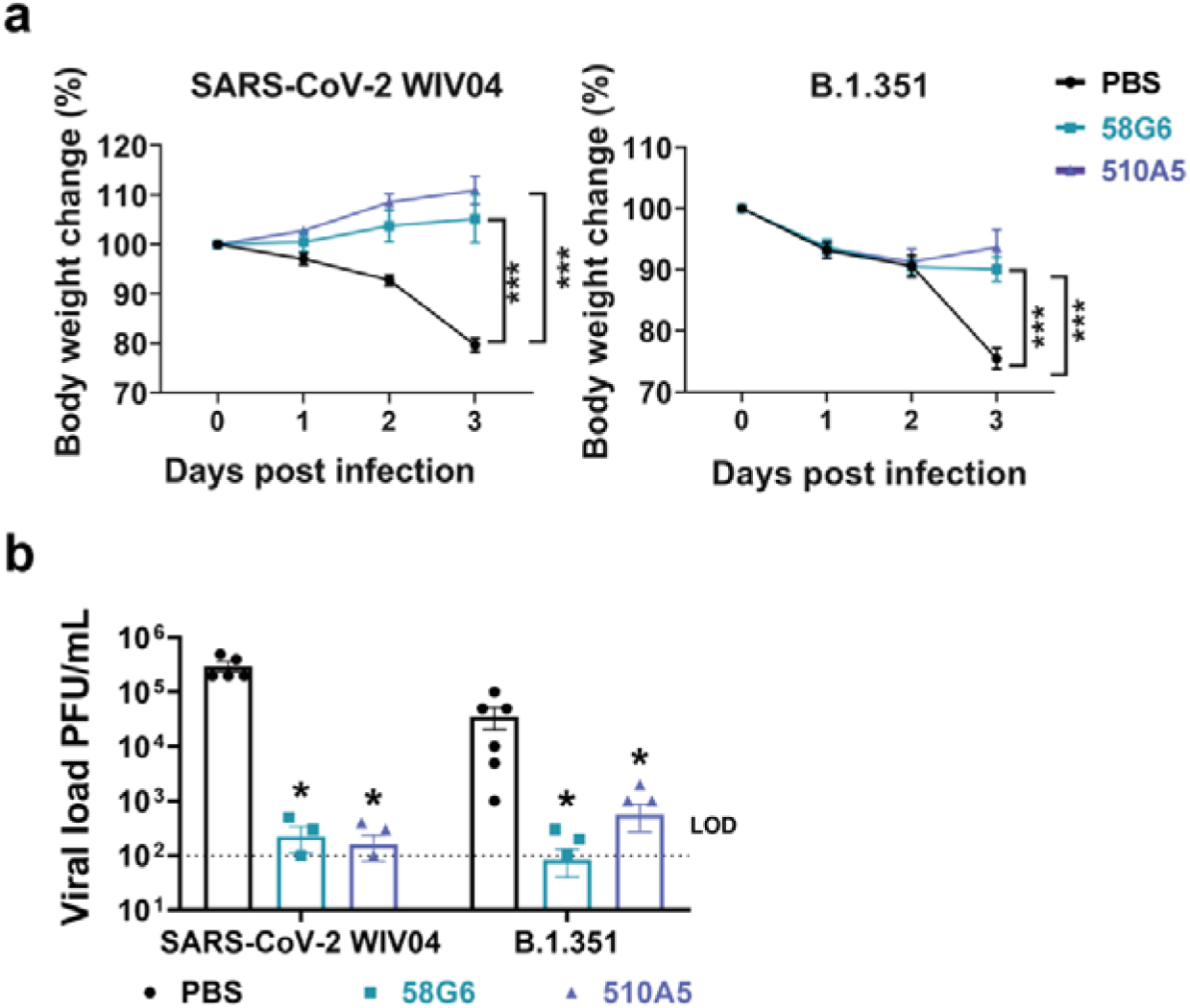
The prophylactic efficacy of 58G6 or 510A5 in hACE2 transgenic mice challenged with authentic viruses. (a) Body weight changes were recorded for PBS (SARS-CoV-2 (WIV04): n = 5; B.1.351: n = 6), 58G6 (SARS-CoV-2 (WIV04): n = 4; B.1.351: n = 7) and 510A5 (SARS-CoV-2 (WIV04): n = 5; B.1.351: n = 7) treatment groups. All the mice received one dose of antibodies (10 mg/kg body weight) injected (i.p.) 24 hours prior to the intranasal challenge with SARS-CoV-2 (WIV04) (left) or B.1.351 (right). Equal volume of PBS was used as negative control. The weight loss was recorded over 3 days. (b) The virus loads in infected lungs were determined by PRNT at 3 days post infection (dpi). \**p* < 0.1, \*\*\**p* < 0.001.

## Discussion

The persistence of COVID-19 has led to generation of mutational variants and immunological adaptation of SARS-CoV-2^1–11^. Newly emerged B.1.351 in South Africa has been reported to confer resistance to neutralization from multiple available mAbs, convalescent plasma and vaccinee sera, posting a high re-infection risk^2–9^. In the present study, we identified 20 neutralizing Abs with high potency against authentic SARS-CoV-2 virus, from a RBD specific mAb reservoir. Among them, 58G6 and 510A5 exhibit high neutralizing capabilities against the authentic virus. Remarkably, these 2 mAbs can efficiently neutralize authentic B.1.351 virus, comparable to most effective neutralizing Abs reported up to date^2,5,6,9^. Their IC_50_s against this variant were as low as approximately 2 ng/mL, hence we termed these 2 mAbs as ultrapotent neutralizing Abs. Such profound neutralizing potencies were confirmed in vivo where the prophylactic treatment of these 2 mAbs could efficiently protect the transgenic mice carrying hACE2 against the airway exposure of authentic SARS-CoV-2 and B.1.351 viruses. These results put 58G6 and 510A5 at the center stage for the development of clinically effective therapeutic regiments against the current COVID-19 pandemic.

In order to understand the high neutralizing potency of our mAbs against SARS-CoV-2, we assessed the antigenic landscape of SARS-CoV-2 RBD. We found that all our RBD targeting mAbs could be categorized into 5 groups according to their recognition on the RBD. Interestingly, the epitopes recognized by the majority of our potent neutralizing Abs overlapped with 13G9e, suggesting that it represented one of the vulnerable sites on SARS-CoV-2 RBD. The other 4 of the top 20 mAbs competed with 510A5 for the binding of RBD at 510A5e. It is worth mentioning that these 2 regions may correspond to 2 separate classes of epitopes recognized by the largest numbers of RBD specific neutralizing Abs, as described in recent studies (Extended Data Fig. 7 and 12)^18,27^.

In detail, we identified that 58G6 recognized a region consisted of amino acids 450-458 in the RBD. Of note, recent cryo-EM structure analysis has revealed 3 key ACE2-interacting residues (S^Y453^, S^L455^, and S^F456^)^13,14^, indicating that S^450-458^ may be the critical site taken into consideration for SARS-CoV-2 prophylaxis. we found at least one specific hydrogen bond within this region, between 58G6 and RBD, that may contribute to recognition of the unique linear region by 58G6, rather than 13G9. Although certain steric proximity of 13G9 to S^450-458^ has been observed, it needs to be pointed out that no specific linear binding sites have been identified for this mAb.

Moreover, 13G9 and 58G6 both recognized the steric epitope of S^470-495^ on the RBD, which was the key region shared by ACE2 and several reported potent neutralizing Abs against SARS-CoV-2^13,14,22,27^. The cryo-EM analysis revealed a hydrogen bond between N94 in 58G6 and the carbonyl group on the main chain, rather than the side chain, of S^E484^. Common mutation within this region found in current variants, such as S^E484K^ in B.1.351 or P.1 emerged in Brazil^9,11^, may not have significant impact on the affinity of 58G6 to the RBDs of these variants. Indeed, the sustained affinity of 58G6 to B.1.351 S1 has been confirmed by the SPR, which may explain for the potentially broad neutralizing spectrum of 58G6. However, for 13G9, the S^E484K^ mutation in B.1.351 or P.1 may introduce an additional positive charge around R94 within its CDRL3, which may lead to strong electrostatic repulsions between the two residues. This may explain the decreased affinity of 13G9 to B.1.351, hence the slight decrease of neutralizing potency against this variant. As far as we know, we are the first to report that an ultrapotent neutralizing Ab to SARS-CoV-2 with direct contact to S^E484K^ still exhibits exceptional potency against authentic B.1.351 virus.

For the potent 13G9 or 58G6, we noted that the RBDs interacting with the 3 Fabs of Abs are universally in the ‘up’ state. As previously described, such full occupancy in each complex could render RBD completely inaccessible for ACE2^15,18,20,22^. However, the significance of this observed phenomena with 3-“up” conformation in all particles of the Fab-S complex, in another word, its correlation to the neutralization advantages, remains unknown.

Interestingly, we noted that 13G9 and 58G6, though originally isolated from the samples of different COVID-19 convalescent donors, were both transcribed from *IGHV1-58* and *IGKV3-20*. These 2 variant regions were also genetically responsible for a panel of reported neutralizing Abs with high potency against SARS-CoV-2 as well as B.1.351^5,6,21,22,27^. These findings highlighted the otherwise overlooked importance for the pairing of the *IGHV1-58* and the *IGKV3-20* germline genes in neutralizing SARS-CoV-2 and its variant.

In conclusion, we present 2 ultrapotent SARS-CoV-2 RBD specific mAbs with exceptional efficacy against B.1.351, for which a significant proportion of reported neutralizing Abs are impaired. Structural analysis of epitopes revealed the potential neutralizing mechanism of neutralizing Abs against B.1.351 carrying the E484K mutation. These broad-spectrum neutralizing Abs could be promising candidates for the prophylaxis and therapeutic interventions of the pandemic of SARS-CoV-2 variants carrying escape mutations.

## Materials and Methods

### Patient information and Isolation of antibodies to SARS-CoV-2

The 74 mAbs analyzed in our manuscript were derived from a total of 39 COVID-19 convalescent blood samples collected within a 2-month window post discharge. These 39 convalescent patients have an average age of 45 years old, and majority of them exhibited mild symptoms, as described in the previous study^24^. The original studies to obtain blood samples after written informed consent were previously described and had been approved by the Ethics Board of ChongQing Medical University^24^. Briefly, we utilized SARS-CoV-2 RBD as bait to sort the antigen-specific memory B cells from the COVID-19 convalescent patients. The IgG heavy and light chains of mAbs genes in these memory B cells were obtained by single cell PCR and transiently transfected into HEK293T cells for the identification of mAbs with capabilities of the neutralization against SARS-CoV-2 pseudovirus. With such rapid screening system, we were capable to obtain the defined neutralizing Abs within 6 days.

### Recombinant antibody production and purification

A pair of plasmids separately expressing the heavy- and the light-chain of antibodies were transiently co-transfected into Expi293^™^ cells (Catalog No. A14528, ThermoFisher) with ExpiFectamine^™^ 293 Reagent. Then the cells were cultured in shaker incubator at 120 rpm and 8% CO_2_ at 37°C. After 7 days, the supernatants with the secretion of antibodies were collected and captured by protein G Sepharose (GE Healthcare). The bound antibodies on the Sepharose were eluted and dialyzed into phosphate-buffered saline (PBS). The purified antibodies were used in following binding and neutralization analyses.

### Authentic SARS-CoV-2 neutralization assay

The neutralizing potency of mAbs against authentic SARS-CoV-2 virus quantified via qRT-PCR was performed in a biosafety level 3 laboratory of Fudan University. Serially diluted mAbs or mAbs mixture (1:1 with same quality) were incubated with authentic SARS-CoV-2 virus (nCoV-SH01, GenBank: MT121215.1, 100 TCID50) for 1 h at 37 °C. After the incubation, the mixtures were then transferred into 96-well plates, which were seeded with Vero E6 cells. The plates were kept at 37 °C for 48 hrs. And the supernatant viral RNA load of each well was quantified by qRT-PCR. For qRT-PCR, the viral RNA was extracted from the collected supernatant using Trizol LS (Invitrogen) and used as templates for the qRT-PCR analysis by Verso 1-Step qRT-PCR Kit (Thermo Scientific) following the manufacturer’s instructions. PCR primers targeting SARS-CoV-2 N gene (nt 608-706) were as followed, forward: 5’-GGGGAACTTCTCCTGCTAGAAT-3’, and reverse: 5’-CAGACATTTTGCTCTCAAGCTG-3’. qRT-PCR was performed using the LightCycler 480 II PCR System (Roche) with the following program: 50 °C 15 mins; 95 °C 15 mins; 40 cycles of 95 °C 15 seconds, 50 °C 30 seconds, 72 °C 30 seconds. The IC_50_ and IC_80_ of the evaluated mAbs was and calculated by a four-parameter logistic regression using GraphPad Prism 8.0.

The neutralizing potency of mAbs against authentic SARS-CoV-2 and B.1.351 viruses was performed quantified via PRNT in a biosafety level 3 laboratory of Wuhan Institute of Virology. Each mAb sample was serially diluted with DMEM as two folds and the sample quality, mixed with equal volume of authentic SARS-CoV-2 virus (WIV04, GenBank: MN996528.1) or SARS-CoV-2 South Africa strain B.1.351 (NPRC 2.062100001, GenBank: MW789246.1) and incubated at 37 °C for 1 h. Vero E6 cells in 24-well plates were inoculated with the sera-virus mixture at 37 °C; 1 h. Later, the mixture was replaced with DMEM containing 2.5% FBS and 0.8% carboxymethylcellulose. The plates were fixed with 8% paraformaldehyde and stained with 0.5% crystal violet 4 days later. All samples were tested in duplicate and neutralization titers were defined as the serum dilution resulting in a plaque reduction of at least 50%^30^.

### Sequence analysis of antigen-specific mAbs

IMGT/V-QUEST (http://www.imgt.org/IMGT_vquest/vquest) and IgBLAST (https://www.ncbi.nlm.nih.gov/igblast/), MIXCR (https://mixcr.readthedocs.io/en/master/) and VDJtools (https://vdjtools-doc.readthedocs.io/en/master/overlap.html) tools were used to do the variable region analysis and annotation for each antibody clone.

### Production of pseudovirus bearing S protein

pVSVG expressing SARS-CoV-2 S protein was constructed as previously described^29^. The packaging plasmid (VSV-G pseudotyped ΔG-luciferase) encoding either the S protein of SARS-CoV-2, B.1.1.7 or chimeric construct including B.1.351 RBD and S^D614G^ was generated. HEK293T cells were grown to 80% confluency before transfection with VSV-G pseudotyped ΔG-luciferase, pWPXL and pSPAX2. These cells were cultured overnight at 37 °C with 5% CO_2_. DMEM supplemented with 5% fetal bovine serum and 100 IU/mL of penicillin and 100 μg/mL of streptomycin was added to the inoculated cells, which were cultured overnight for 72 hrs. The supernatant was harvested, filtered by 0.45 μm filter and centrifugated at 300 g for 10 mins to collect the supernatant, then aliquoted and storied at −80 °C.

### Pseudovirus neutralization assay

Serially diluted mAbs with volume of 50 μL were incubated with the same volume of the HEK293T cell supernatants containing the pseudovirus for 1 h at 37 °C. These pseudovirus-antibody mixtures were added to ACE2 expressing HEK293T cells (HEK293T/ACE2). After 72 hrs, the luciferase activities of infected HEK293T/ACE2 cells were detected by the Bright-Luciferase Reporter Assay System (Promega, E2650). The IC_50_ of the evaluated mAbs was tested by the Varioskan LUX Microplate Spectrophotometer (Thermo Fisher), and calculated by a four-parameter logistic regression using GraphPad Prism 8.0.

### Protein expression and purification

To express the prefusion S ectodomain, the gene encoding residues 1-1208 of SARS-CoV-2 S (GenBank: MN908947.3) with a C-terminal T4 fibritin trimerization motif, an HRV-3C protease cleavage site, a Twin-Strep-tag and an 8 × His-tag was synthesized, and cloned into the mammalian expression vector pcDNA3.1, which was a kind gift from L. Sun at Fudan University, China. The gene of the S protein was constructed with proline substitutions at residues 986 and 987, a “GSAS” instead of “RRAR” at the furin cleavage site (residues 682-685) according to Jason S. McLellan’s research^25^.

Expi293 cells (Thermo Fisher Scientific, USA) cultured in Freestyle 293 Expression Medium (Thermo Fisher Scientific, USA) were maintained at 37 °C. Cells were diluted to a density of 2.5 × 10^6^ to 3 × 10^6^ cells per mL before transfection. For protein production, 1.2 mg DNA was mixed with 3 mg polyethyleneimine in 30 mL Freestyle 293 Expression Medium, incubated for 20 mins, then added to 1000 mL of cells^31^. Transfected cells were cultured at 35 °C, and the cell culture supernatant was collected at day 4 to day 5.

The protein was purified from filtered cell supernatants using Strep-Tactin resin (IBA) before being subjected to additional purification by gel filtration chromatography using a Superose 6 10/300 column (GE Healthcare, USA) in 1 × PBS, pH 7.4 (Extended Data Fig. 10a, b).

### Cryo-EM sample preparation and data collection

Purified SARS-CoV-2 S was diluted to a concentration of 1.5 mg/mL in PBS, pH 7.4. 5 μL of purified SARS-CoV-2 S was mixed with 1 μL of 58G6 Fab fragments at 2 mg/mL in PBS and incubated for 30 mins on ice. A 3 μL aliquot of the mixture (added with 0.01% DDM) was applied onto an H_2_/O_2_ glow-discharged, 300-mesh Quantifoil R1.2/1.3 grid (Quantifoil, Micro Tools GmbH, Germany). The grid was then blotted for 3.0 s with a blot force of −1 at 8 °C and 100% humidity and plunge-frozen in liquid ethane using a Vitrobot (Thermo Fisher Scientific, USA). Cryo-EM data sets were collected at a 300 kV Titan Krios microscope (Thermo Fisher Scientific, USA) equipped with a K3 detector (Gatan, USA). The exposure time was set to 2.4 s with a total accumulated dose of 60 electrons per Å^2^, which yields a final pixel size of 0.82 Å. 2605 micrographs were collected in a single session with a defocus range comprised between 1.0 and 2.8 μm using SerialEM. The sample preparation and data collection for the SARS-CoV-2 S-13G9 Fab complex were in accordance with the SARS-CoV-2 S-58G6 Fab complex. The statistics of cryo-EM data collection can be found in Extended Data Table 1.

### Cryo-EM data processing

All dose-fractioned images were motion-corrected and dose-weighted by MotionCorr2 software^32^ and their contrast transfer functions were estimated by cryoSPARC patch CTF estimation^33^. For the dataset of SARS-CoV-2 S-58G6 Fab complex, a total of 1,255,599 particles were auto-picked using the template picker and 820,872 raw particles were extracted with a box size of 512 pixels in cryoSPARC^33^. The following 2D, 3D classifications, and refinements were all performed in cryoSPARC. 237,062 particles were selected after two rounds of 2D classification, and these particles were used to do Ab-Initio reconstruction in six classes. Then these six classes were used as 3D volume templates for heterogeneous refinement with all selected particles, with 108,020 particles converged into the SARS-CoV-2 S-58G6 Fab class. Next, this particle set was used to perform non-uniform refinement, yielding a resolution of 3.56 Å.

For the dataset of SARS-CoV-2 S-13G9 Fab complex, a total of 445,137 particles were auto-picked using the template picker and 266,357 raw particles were extracted with a box size of 512 pixels in cryoSPARC. The following 2D, 3D classifications, and refinements were all performed in room temperature (RT). 70,519 particles were selected after two rounds of 2D classification, and these particles were used to do Ab-Initio reconstruction in six classes. Then these 6 classes were used as 3D volume templates for heterogeneous refinement with all selected particles, with 52,880 particles converged into the SARS-CoV-2 S-13G9 Fab class. Next, this particle set was used to perform non-uniform refinement, yielding a resolution of 3.92 Å.

Although the overall resolution for these structures is up to 3.5 Å - 3.6 Å for 58G6 and 3.9 Å - 4.0 Å for 13G9, the maps for the binding interface between RBD and Fabs are quite weak due to the conformational heterogeneity of the RBD, which is similar to previous structural investigations^15,18,22,34^. To improve the resolution for the binding interface, we subsequently added local refinement processing. A local reconstruction focusing on the RBD-Fabs region was carried out. Furthermore, the density map for the binding interface could be improved further by local averaging of the RBD-Fab equivalent copies, finally yielding a 3.5 Å map of the region corresponding to the 58G6 variable domains and the RBD (Extended Data Fig. 10c, f). Similarly, we improve the local resolution between the 13G9 variable domains and the RBD up to 3.8 Å (Extended Fata Fig. 10g, j).

Local resolution estimation, filtering, and sharpening were also carried out using cryoSPARC. The full cryo-EM data processing workflow is described in Extended Data Fig. 10 and the model refinement statistics can be found in Extended Data Table 1.

### Model Building and Refinement

To build the structures of the SARS-CoV-2 S-58G6 Fab and S-13G9 Fab complexes, the structure of the SARS-CoV-2 S glycoprotein in complex with the C105 neutralizing antibody Fab fragment^15^ (PDB: 6XCN) was placed and rigid-body fitted into the cryo-EM electron density maps using UCSF Chimera^35^, respectively. Both of the 58G6 and 13G9 Fab models were first predicted using Phyre2^26^ and then manually built in Coot 0.9^36^ with the guidance of the cryo-EM electron density maps, and overall real-space refinements were performed using Phenix 1.18^37^. The data validation statistics are shown in Extended Data Table 1.

### Creation of Figures

Figures of molecular structures were generated using PyMOL^38^ and UCSF ChimeraX^39^.

### The antibody binding kinetics and the competition with ACE2 measured by SPR

The affinity of the neutralizing Abs binding to the S1 subunit of SARS-CoV-2 or B.1.351 was measured using the Biacore X100 platform at RT. A CM5 chip (GE Healthcare) was linked with anti-human IgG-Fc antibody to capture about 9000 response units of the neutralizing Abs. The gradient concentrations of SARS-CoV-2 S1 or an artificial chimeric construct carrying 3 mutations on B.1.351 RBD and S^D614G^ (B.1.351 S1) (Sino Biological, Beijing, China) were prepared (2-fold dilutions, from 50 nM to 0.78 nM) with HBS-EP^+^ Buffer (0.01 M HEPES, 0.15 M NaCl, 0.003 M EDTA and 0.05% (v/v) Surfactant P20, pH 7.4), and sequentially injected into the chip and monitored for the binding kinetics. After the final reading, the sensor surface of the chip was regenerated with 3 M MgCl2 (GE) before the measurement of the next mAb. The affinity was calculated with Biacore X100 Evaluation Software (Version:2.0.2) using 1:1 binding fit model.

To determine competition with the ACE2 peptidase domain, SARS-CoV-2 RBD was coated on a CM5 sensor chip via amine group for a final RU around 250. The top 20 neutralizing Abs (20 μg/mL) were injected onto the chip until binding steady-state was reached. ACE2 (20 μg/mL) was then injected for 60 seconds. Blocking efficacy was determined by comparison of response units with and without prior antibody incubation.

### Competitive ELISA

For competitive ELISA used in epitope mapping of mAbs, 2 μg/mL recombinant RBD-his (Sino Biological, Beijing, China) was added in 384-well plates and incubated at 4 °C overnight. 50 μg/mL mAbs per well were added. The plates were incubated at 37 °C for 1 h and then washed. Biotinylation of mAbs (the top 20 neutralizing Abs and 81A11, previously reported SARS-CoV CR3022^28^) was performed using the EZ-link NHS-PEO Solid Phase Biotinylation Kit (Pierce) according to the manufacturer’s protocol and purified using MINI Dialysis Unit (ThermoFisher, 69576). 500 ng/mL biotinylated mAbs were added to each well, and the plates were incubated at 37 °C for 1 h. ALP-conjugated streptavidin (Mabtech, Sweden, 3310-10) was added at 1:1000, followed by an incubation of 30 mins at 37 °C. For the quantification of bound IgG, PNPP (Thermo Fisher) was added at 1 mg/mL and the absorbance at 405 nm was measured by the MultiSkan GO fluoro-microplate reader (Thermo Fisher).

### Western blot analysis

The recombinant RBD protein was mixed with 5 × loading buffer (Beyotime, Shanghai, China) and denatured for 5 mins at 100 °C. The denatured proteins (200 ng) were subjected to electrophoresis with 10% SDS-polyacrylamide gel and then transferred to PVDF membranes. After blocking by skim milk (Biofroxx), the membranes were incubated at 4 °C overnight, with the purified mAbs as primary Abs. The next days, the membranes were washed with TBST and incubated with HRP-conjugated Goat-anti-human Fc antibody (Abcam, ab99759, 1:10000) for 1 h at RT. The membranes were examined on ChemiDoc Imaging System (Bio-rad).

### Peptide ELISA

Peptide ELISA was performed with synthesized peptides overlapping with 5 amino acids (Genescripts, Wuhan, China). These peptides were tethered by N-terminal biotinylated linker peptides (biotin-ahx), except for the first peptide at the N-terminus, whose biotin was linked to the C terminus instead. The RBD9-1 amino acid residues were selected and mutated to alanine and synthesized by Genescripts (Wuhan, China). 50 μL synthesized peptide was added to the streptavidin-coated 384-well plate in duplets to make a final concentration of 5 μg/mL. The plates were incubated for 2 hrs at RT. After washing, the plates were blocked with Protein-Free Blocking Buffer (Pierce, USA, 37573) at RT for 1 h and incubated with 10 μg/mL testing mAbs at RT for another 1 h. Reacted mAbs were detected using ALP-conjugated Goat F(ab’)_2_ Anti-Human IgG (Fab’)_2_ secondary antibody conjugated with ALP (Abcam, ab98532, 1:2000) for 30 mins at RT, followed with quantification detection.

For the ACE2 competitive peptide ELISA, 5 μg/mL synthesized RBD9-1 was immobilized on the streptavidin-coated 384-well plate at RT for 2 hrs. After washing with Protein-Free Blocking Buffer, the plates were blocked with this blocking buffer. Next, serial diluted 58G6 (20-0.625 μg/mL) in 50 μL of the blocking buffer were added into plate and the plates were incubated at RT for 1 h. Then, the plate incubated with 2 μg/mL ACE2 at RT for another 1 h. The ELISA plates were washed 4 times by blocking buffer and 50 μL Goat F(ab’)_2_ Anti-Human IgG (Fab’)_2_ secondary antibody conjugated with ALP (Abcam, ab98532, 1:2000) was incubated with the plate at RT for 30 mins. The plate was washed and followed with quantification detection.

### Authentic SARS-CoV-2 and B.1.351 viruses and animal study

Authentic SARS-CoV-2 (WIV04) and B.1.351 (NPRC 2.062100001) viruses were propagated on the Vero-E6 cells and titrated by single layer plaque assay with standard procedure. The hACE2 mouse model was used to evaluate the efficacy of 58G6 and 510A5 monoclonal antibodies *in vivo*. Six-to eight-week-old female hACE2 mice were treated with 58G6 or 510A5 monoclonal antibody at a concentration of 10 mg/kg by intraperitoneal route, respectively. The mice treated with PBS were used as the negative control. 24 hours later, all mice were intranasally infected with 10^5^ PFU authentic SARS-CoV-2 or B.1.351 viruses in a total volume of 50 μL. At 3 days post infection of SARS-CoV-2 or B.1.351, the lungs of mice were collected for viral load determination using plaque assay^40^.

### Data analysis

Data are shown as mean ± SEM. Two-group comparisons were performed by Student’s t-test. The difference was considered significant if p < 0.05.

## Data availability

The coordinates and structure factor files for the 13G9/SARS-CoV-2 RBD complex and 58G6/SARS-CoV-2 RBD complex have been deposited in the Protein Data Bank (PDB) under accession number 7E3K and 7E3L respectively.

## Acknowledgments

This study was supported by the Emergency Project from Chongqing Medical University and Chongqing Medical University fund (X4457) with the donation from Mr. Yuling Feng. We acknowledge the clinical laboratories of Yongchuan Hospital of Chongqing Medical University and the Third Affiliated Hospital of Chongqing Medical University for providing blood samples. We are grateful to Xiaoxiao Gao and Cheng Peng for their technical help, and to Wuhan National Biosafety Laboratory running team, including engineer, biosafety, biosecurity, and administrative staff. The SARS-CoV-2 South Africa strain (NPRC 2.062100001) was provided by Guangdong Provicial Center for Disease Control and Prevention. We also thank all healthy individuals participated in this study.

## Author contributions

A.J. and A.H. conceived and designed the study. F.L. and H.J. were responsible for antibody production and purification. J.W., K.W., J.H., S.L., N.T., G.Z. and Q.G. conducted the pseudovirus neutralization assays and Y.X., C.G., Y.W., W.X., X.C., D.Q. and Z.Y. performed authentic SARS-CoV-2 neutralization assays. S.L. and Y.H. played an import role in data analysis of neutralizing Abs sequences. T.L., Y.W., Y.L., S.S., Q.C., F.G. and M.S. performed ELISA, competitive ELISA and peptide ELISA. X.H., C.H., R.W. and S.M. were responsible for SPR assay for the affinity of these neutralizing Abs and competition of these neutralizing Abs with ACE2. H.G., F.L., Y.G., W.W., X.J. and H.Y. carried out the cryo-EM studies. H.Z, Y.Z, Z.Z, H.Z, N.L and B.Z were responsible for the prophylactic test of neutralizing Abs for hACE2 mice challenged with SARS-CoV-2 and B.1.351. L.L. and C.H. generated figures and tables, and take responsibility for the integrity and accuracy of the data presentation. A.J., T.L., W.W. and H.G wrote the manuscript.

## Ethics statement

The project “The application of antibody tests patients infected with SARS-CoV-2” was approved by the ethics committee of Chongqing Medical University. Informed consents were obtained from all participants.

All the mice were cared in accordance with the recommendations of National Institutes of Health Guidelines for the Care and Use of Experimental Animals. Viral infections were conducted in an animal biosafety level 3 (ABSL-3) facility at Wuhan Institute of Virology under a protocol approved by the Laboratory Animal Ethics Committee of Wuhan Institute of Virology, Chinese Academy of Sciences (Permit number: WIVA26201701).

