## Supplemental Figure and Table for "Ultrapotent SARS-CoV-2 neutralizing antibodies with protective efficacy against newly emerged mutational variants"

Includes:

Extended Data Figs. 1-12

Extended Data Tables 1-2

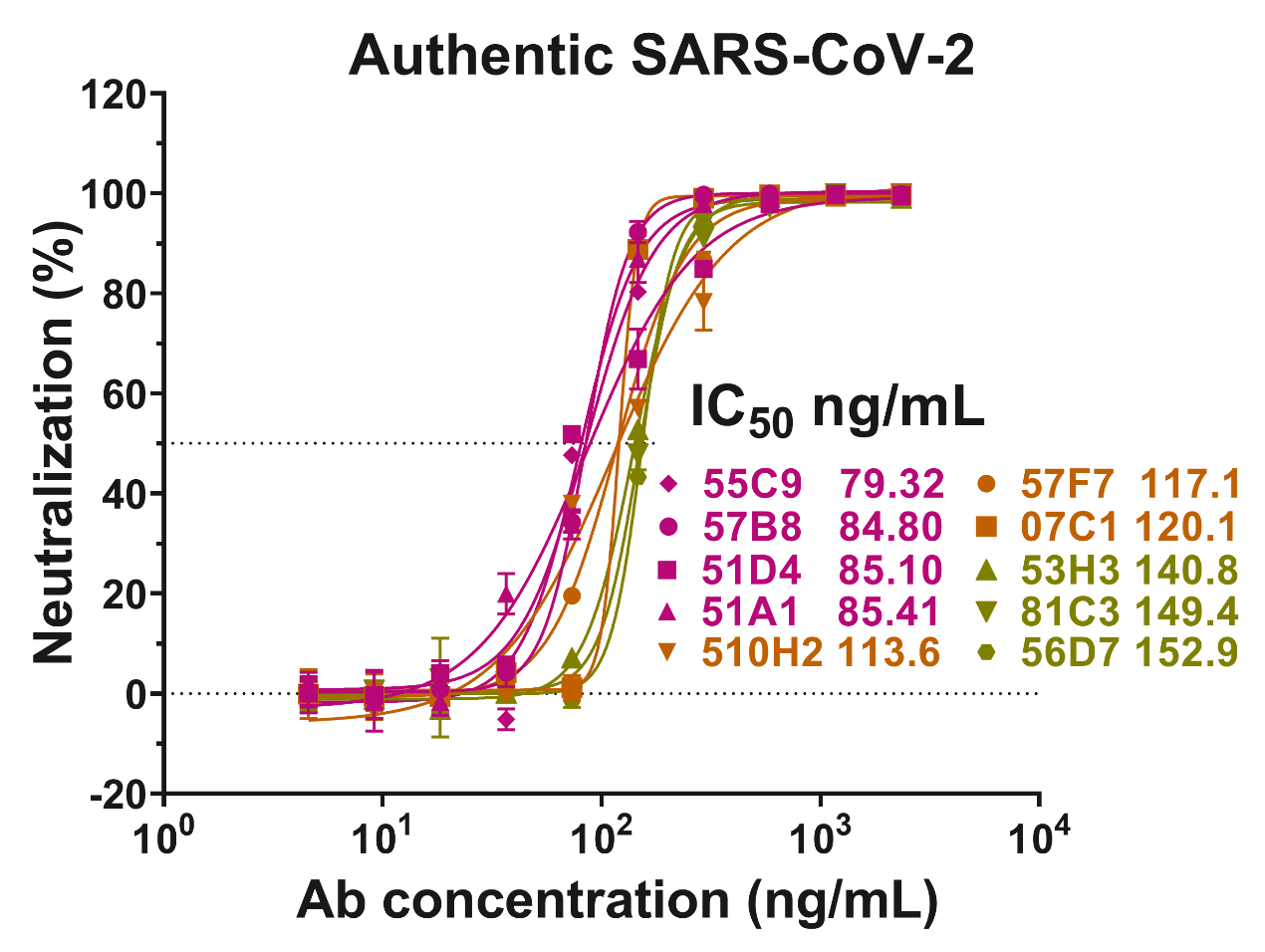

**Extended Data Fig. 1 | The neutralizing efficacy of top 11-20 mAbs against authentic SARS-CoV-2 virus.** The capabilities of the top 11-20 mAbs were measured by authentic SARS-CoV-2 (nCoV-SH01) neutralization assay and quantified by qRT-PCR. Dashed line indicated 0% or 50% reduction in viral neutralization. Data for each mAb were obtained from a representative neutralization experiment of three replicates, presented as mean ± SEM.

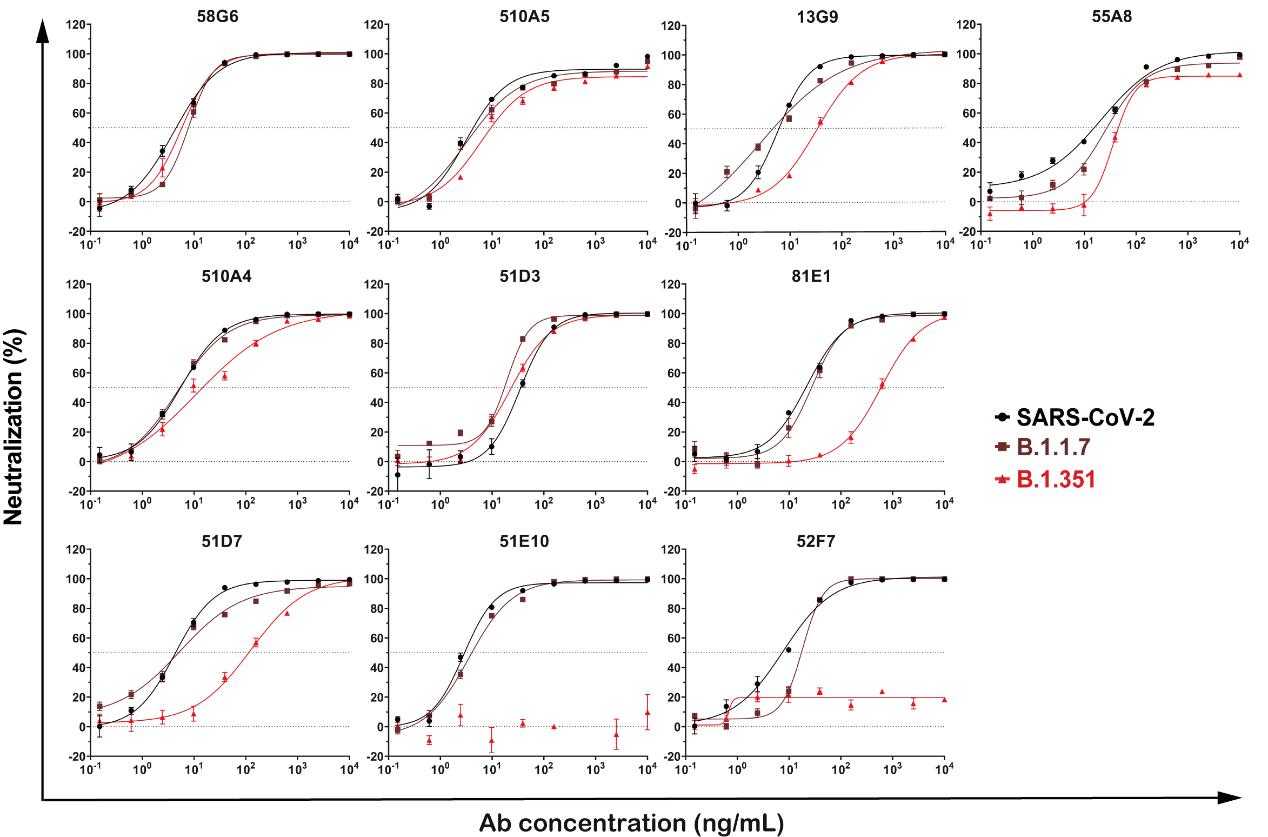

**Extended Data Fig. 2 | The neutralizing capabilities of selected mAbs against SARS-CoV-2, B.1.1.7 and B.1.351 pseudoviruses.** The neutralizing potencies of the top 10 mAbs against SARS-CoV-2, B.1.1.7 and B.1.351 were measured by pseudovirus neutralization assay. Dashed line indicated 0% or 50% reduction in the viral neutralization. Data for each mAb were obtained from a representative neutralization experiment of three replicates, presented as mean ± SEM.

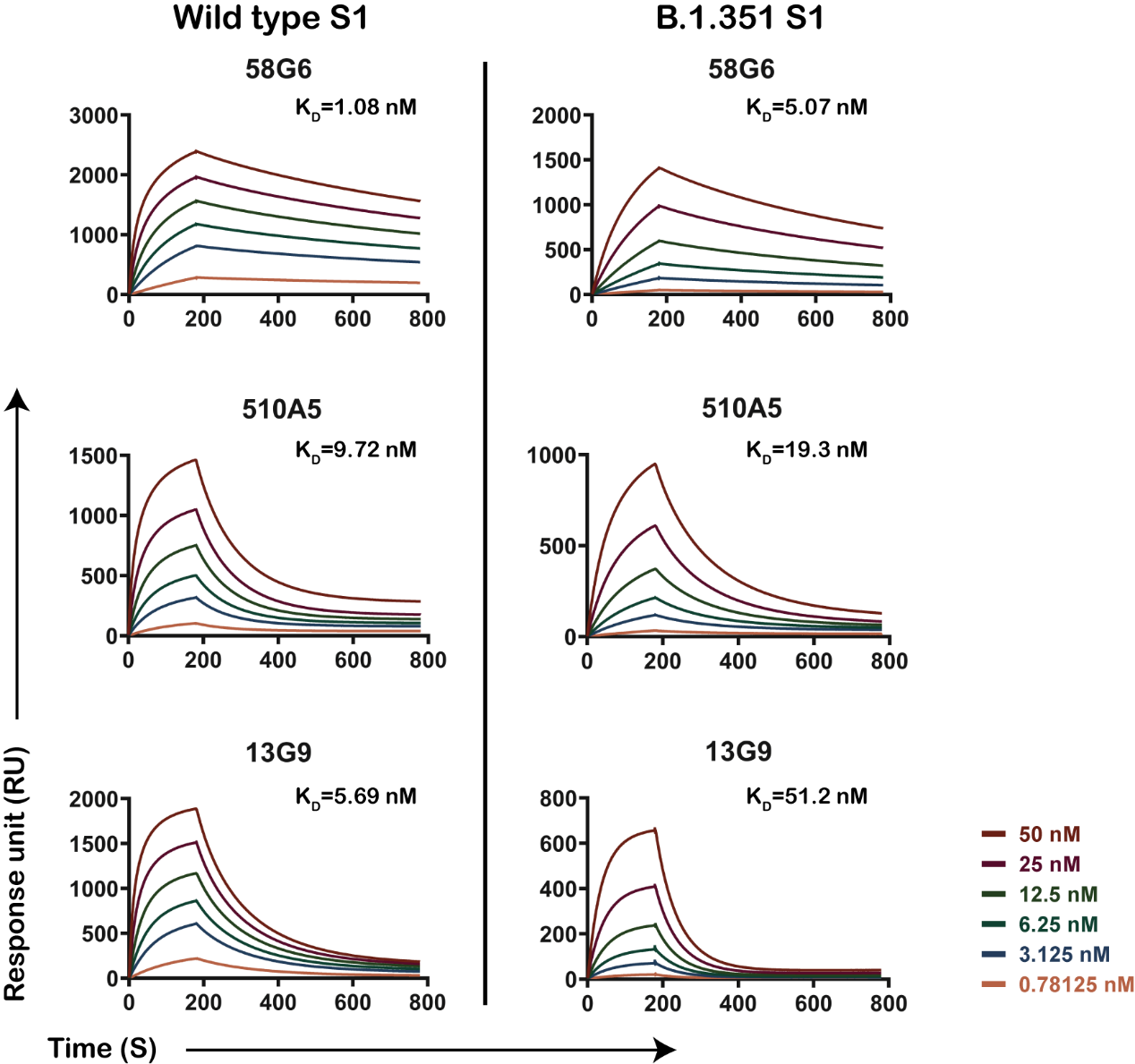

**Extended Data Fig. 3 | The binding kinetics of 58G6, 510A5 and 13G9 to SARS-CoV-2 S1 or B.1.351 S1.** The purified mAbs were coated on the CM5 sensor chip followed by the injection of various concentrations of soluble recombinant SARS-CoV-2 S1 or B.1.351 S1 proteins. Data are representative of at least 2 independent experiments.

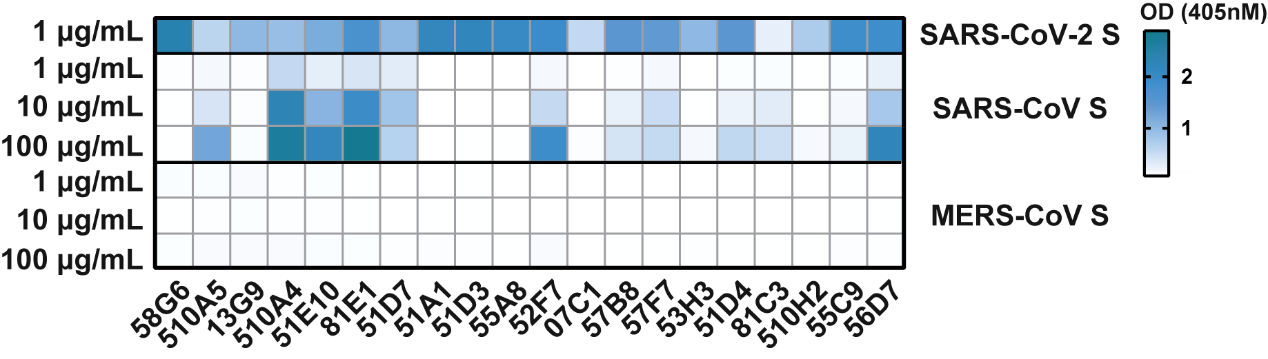

**Extended Data Fig. 4 | The binding of the top 20 mAbs to SARS-CoV-2 S, SARS-CoV S and MERS-CoV-2 S.** The binding of these top 20 mAbs to the S protein of SARS-CoV-2, SARS-CoV or MERS-CoV was tested with various concentrations of the mAbs via ELISA. Data are representative of at least 2 independent experiments performed in technical duplicate.

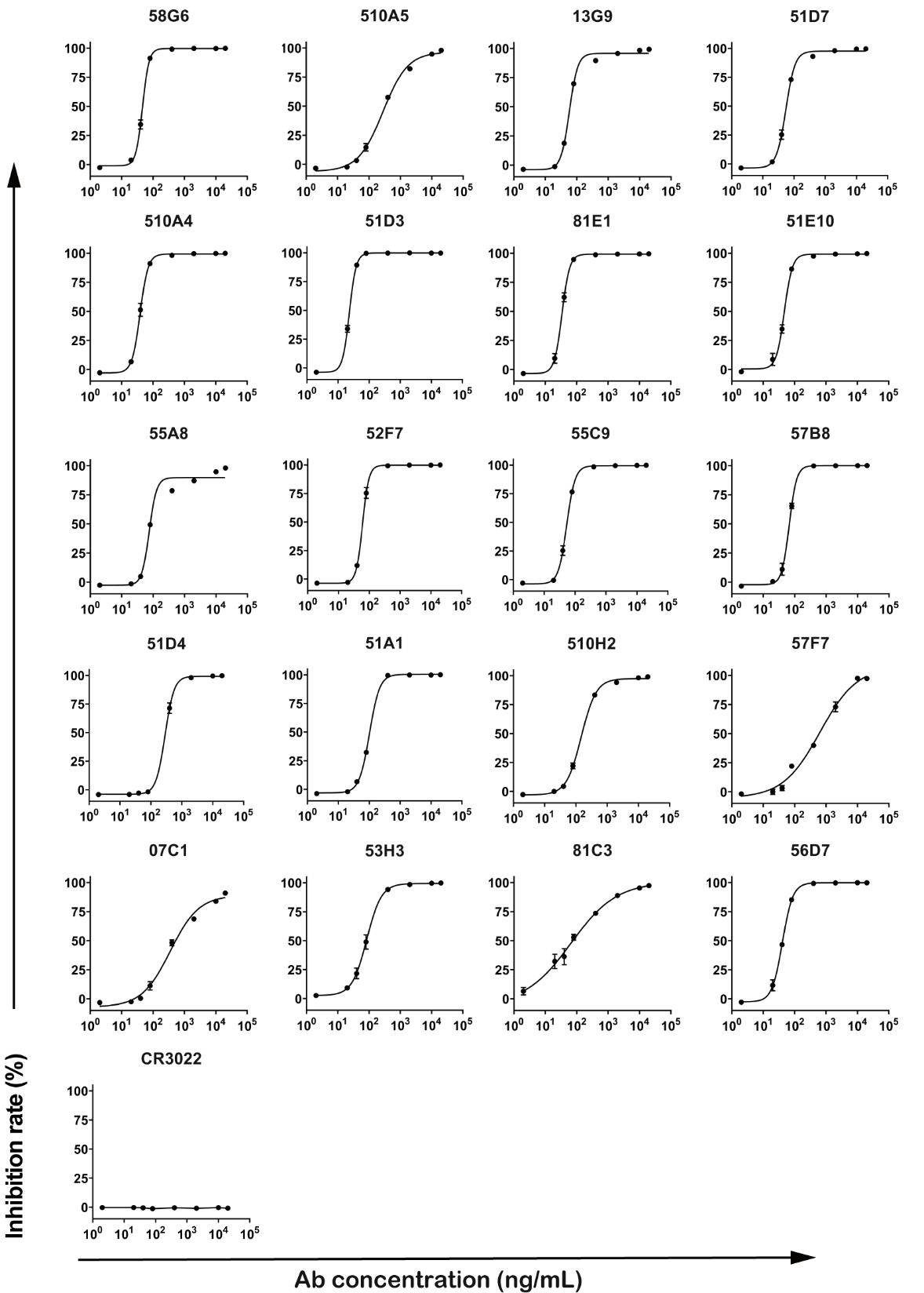

**Extended Data Fig. 5 | The inhibition curves for the neutralizing Abs on the ACE2-RBD interaction.** Competitive ELISA for the neutralizing Abs to inhibit the ACE2-RBD interaction. The SARS-CoV specific mAb CR3022 was used as the negative control. Data are representative of at least 2 independent experiments performed in technical duplicate, presented as mean ± SEM.

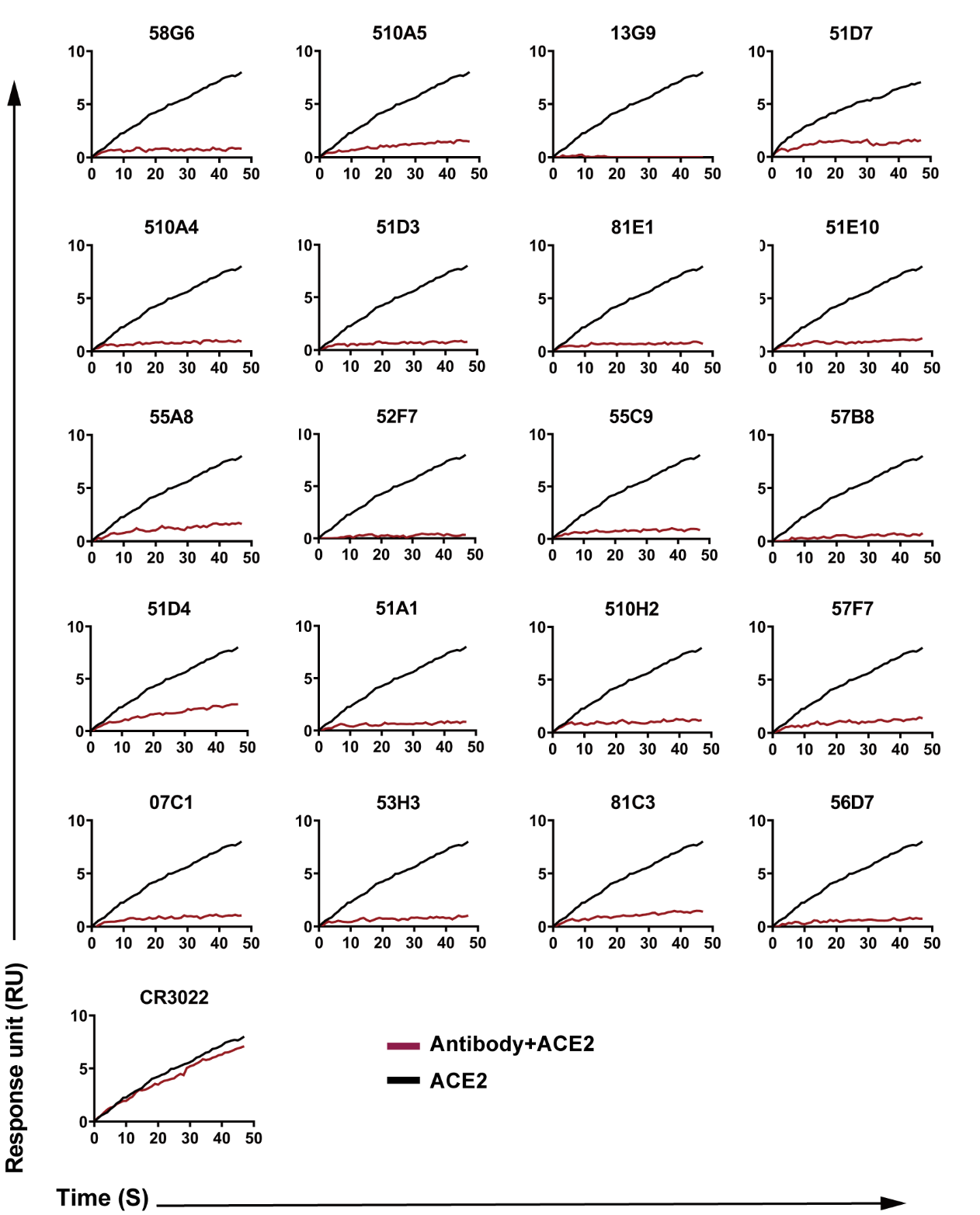

**Extended Data Fig. 6 | Neutralizing Abs competing with ACE2 for the binding of RBD.** In SPR assay, the purified soluble SARS-CoV-2 RBD protein was coated onto a CM5 sensor chip followed by injection of individual neutralizing Abs at concentration of 20 μg/mL. The competition capacity of each antibody was indicated by the level of reduction in the response unit of ACE2 comparing with or without prior antibody incubation. The SARS-CoV RBD mAb CR3022 was used as the negative control.

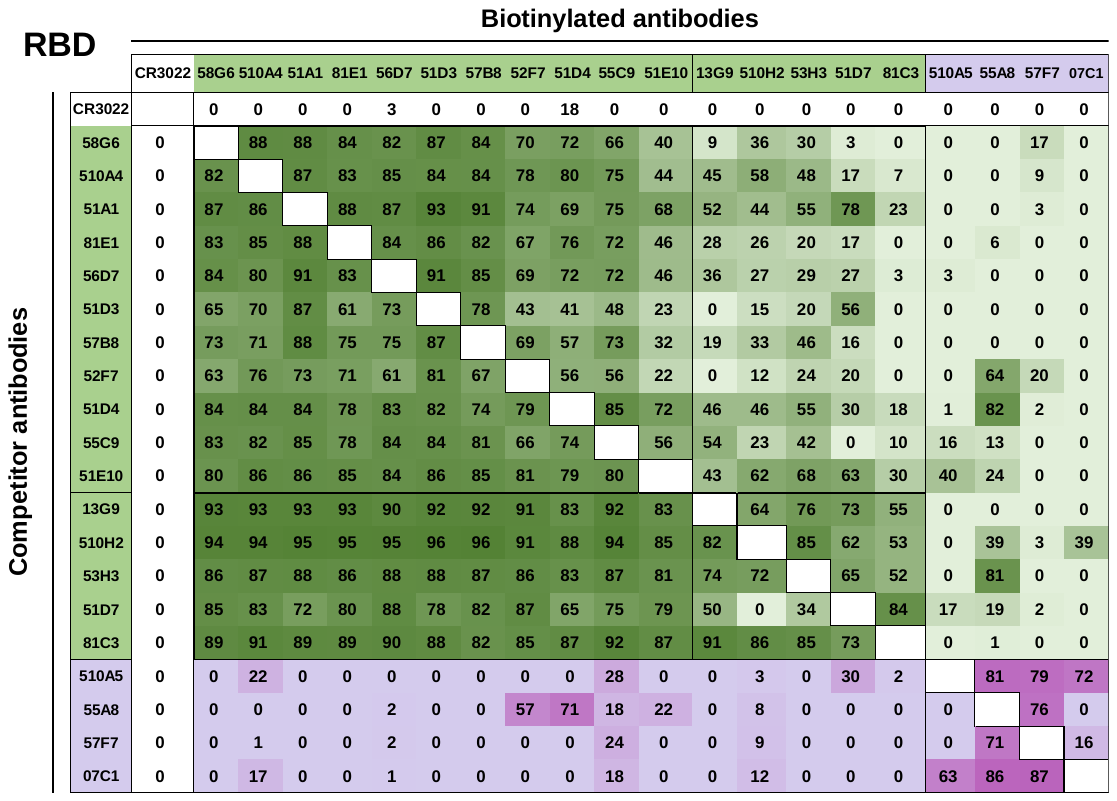

**Extended Data Fig. 7 | The competitive analysis of potential epitopes recognized by the top 20 neutralizing Abs.** Each of our 20 mAb was biotinylated and competed with other unmodified mAbs for the identification of corresponding epitopes. The number in each box indicated the inhibition rate between two mAbs tested by competitive ELISA. Results are representatives of two independent experiments.

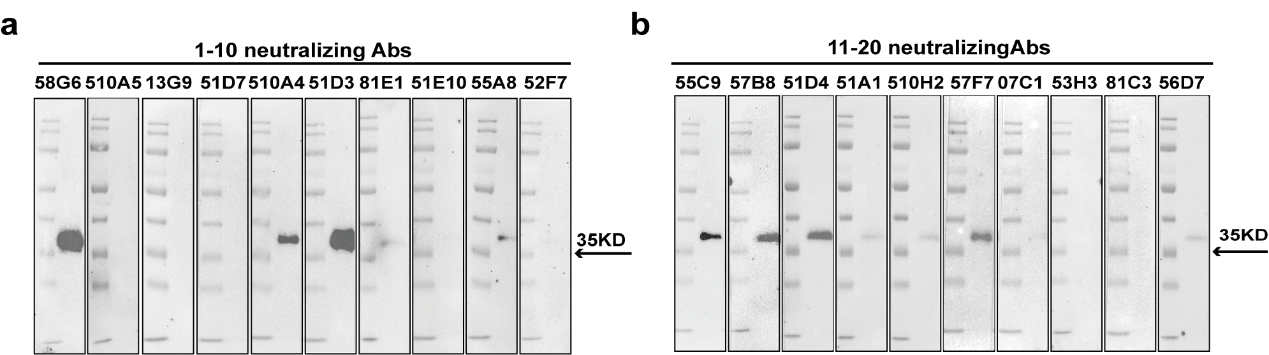

**Extended Data Fig. 8 | The binding ability of purified mAbs to the denatured RBD.** Western blot results of the 1-10 (a) and 11-20 (b) neutralizing Abs binding to the denatured RBD. Data are representative of at least 2 independent experiments performed.

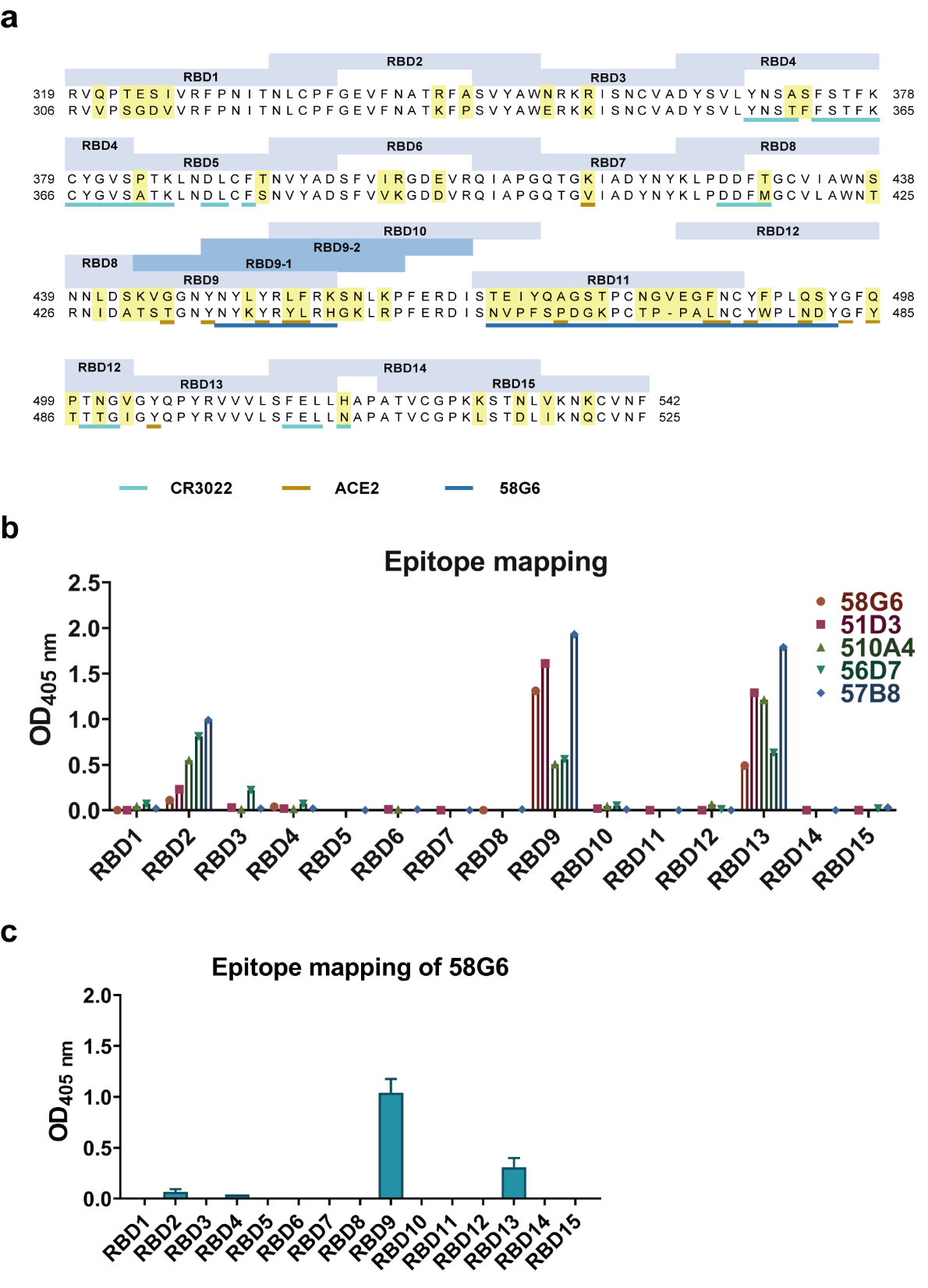

**Extended Data Fig 9. | The binding ability of purified mAbs to the RBD related peptides.** (a) The amino acid sequences for the RBDs of SARS-CoV-2 and SARS-CoV were aligned for comparison. Linear peptides designed for SARS-CoV-2 were shown in ash blue and the names of the peptides were indicated in the boxes above. The non-conserved residues between two viruses were displayed in yellow. The binding sites for CR3022, 58G6 and ACE2 were distinguished by underlines of different colors indicated below. (b) The interactions of 5 out of 9 mAbs (58G6, 510A4, 51D3, 57B8 and 56D7) capable of binding to the denatured RBD to the linear peptides (RBD1-RBD15) were analyzed by peptide ELISA. The other 4 mAbs were uncapable to react with synthesized peptides. (c) The interactions of 58G6 to RBD1-RBD15 were analyzed by peptide ELISA. Data are representative of at least 2 independent experiments performed in technical duplicate. The mean ± SEM of duplicates are shown.

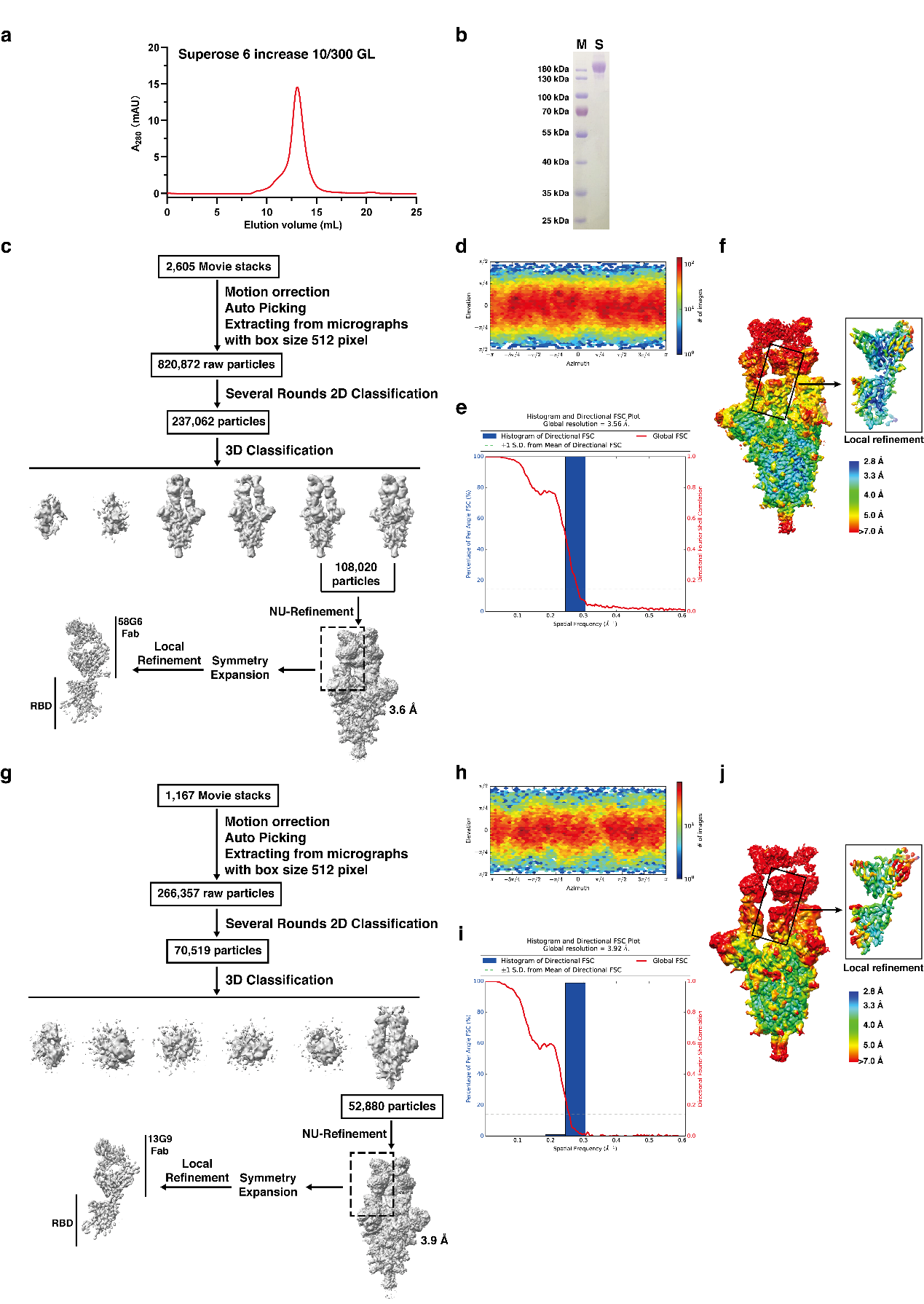

**Extended Data Fig. 10 | Workflow for 58G6 and 13G9 cryo-EM 3D Reconstructions.** (a) Gel filtration profile of the affinity-purified SARS-CoV-2 S trimer. (b) SDS-PAGE analysis of the SARS-CoV-2 S protein. (c, g) Cryo-EM data processing workflow of 58G6 (c) and 13G9 (g). (d, h) The viewing direction distribution plot for SARS-CoV-2 S in complex with 58G6 Fab (d) and 13G9 (h). (e, i) Global FSC and histogram. (f, j) Cryo-EM density of SARS-CoV-2 S-Fab complexes, colored according to local resolution, including locally refined reconstruction of the RBD-58G6 (f) or RBD-13G9 (j) variable domains.

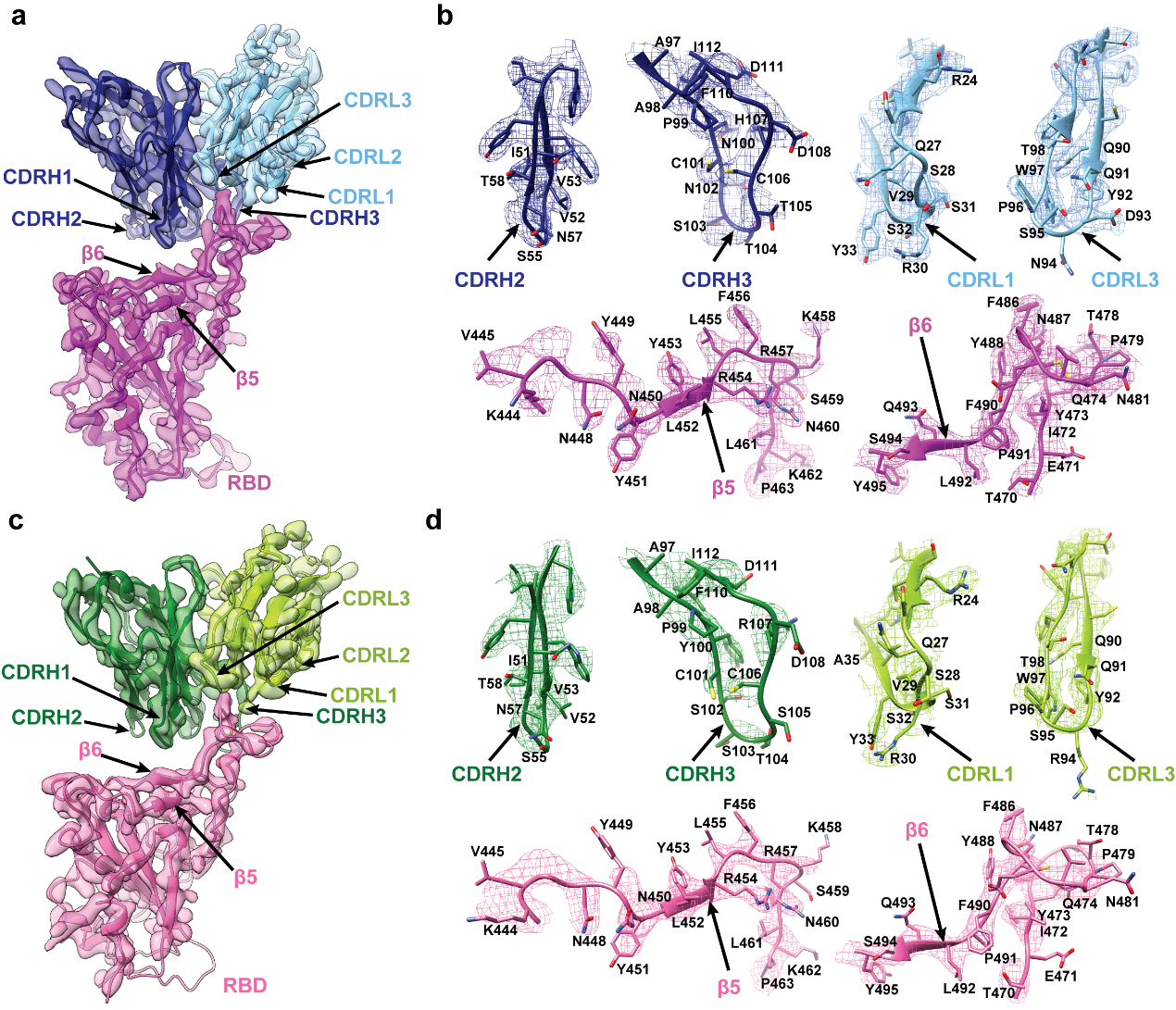

**Extended Data Fig. 11 | Density maps and atomic models.** (a, c) Cryo-EM maps of the binding interface between SARS-CoV-2 RBD and 58G6 Fab (a) or 13G9 Fab (c) variable domains. The color scheme is the same as in Fig. 3. (b, d) Density maps (mesh) and related 58G6 CDRs (b) or 13G9 CDRs (d) atomic models. Residues are shown as sticks, oxygen atoms are colored red, nitrogens are colored blue and sulfurs are shown in yellow.

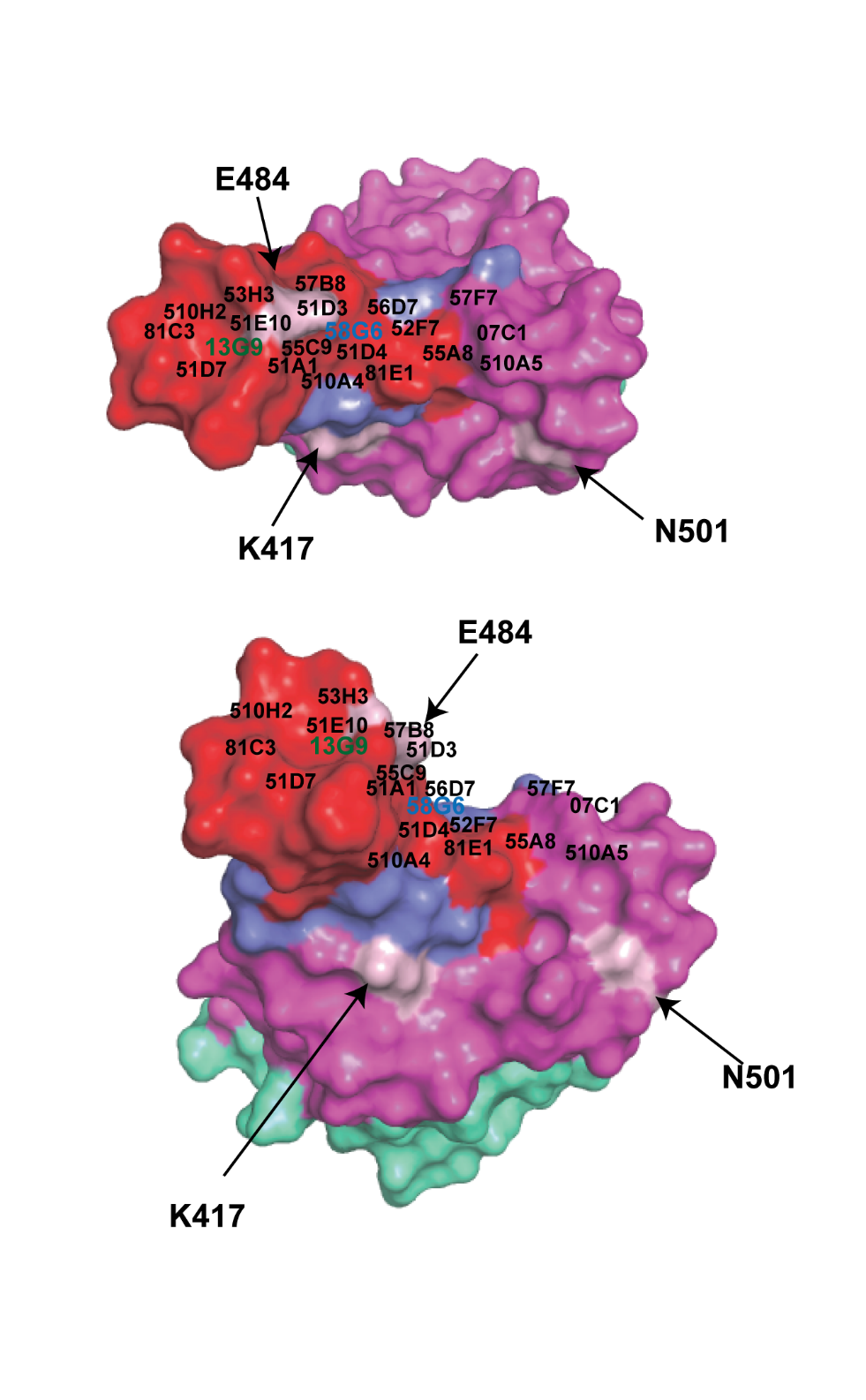

**Extended Data Fig. 12 | Mapping of the epitopes for the top 20 neutralizing Abs onto the surface of the viral RBD.** The data for mapping calculations were from Extended Data Fig. 7. The view is chosen for clarity and is related to that shown in Fig. 5a, b. The epitopes for 58G6 and 13G9 were distinguished according to the color shown in Fig. 5a, b. The epitope for CR3022 was colored cyan. Positions of the S^K417^, S^E484^, and S^N501^ were labeled.

**Extended Data Table 1. Cryo-EM data collection and models refinement statistics**

**EM Data collection and reconstruction statistics**

Protein	S-58G6 Fab	S-13G9 Fab

Voltage (kV)	300	300

Detector	K3	K3

Pixel size (Å)	0.82	0.82

Electron dose (e^-^/ Å^2^)	60	60

Defocus range	1.0-2.8	1.0-2.8

Micrographs collected	2605	1167

Particles initial/final	820,872/106,020	266,357/52,880

Final resolution (Å)	3.56	3.92

**Models refinement and validation statistics**

Protein	S-58G6 Fab	S-13G9 Fab

**RMSD**

Bond lengths (Å)	0.007	0.008

Bond angles (°)	1.129	1.173

**Ramachandran statistics**

Favored (%)	91.00	91.98

Allowed (%)	8.91	7.93

Outliers (%)	0.09	0.09

Rotamer outliers (%)	0.32	0.12

Clash score	8.99	12.66

C-beta outliers (%)	0.10	0.03

CaBLAM outliers (%)	4.01	3.93

**Extended Data Table 2 | The information of 58G6 and 13G9 variable genes and sequences.**

| **mAbs** | **VDJ information** | | | | **CDR information** | | | | | |
| --- | --- | --- | --- | --- | --- | --- | --- | --- | --- | --- |
|  | **VH** | **JH** | **VL** | **JL** | **CDRH1** | **CDRL1** | **CDRH2** | **CDRL2** | **CDRH3** | **CDRL3** |
| 58G6 | IGHV1-58 | IGHJ3-2 | IGKV3-20 | IGKJ1-1 | GFTFSSSA | QSVRSSY | IVVGSGNT | GAS | AAPNCNSTTCHDGFDI | QQYDN*****SPWT |
| 13G9 | IGHV1-58 | IGHJ3-2 | IGKV3-20 | IGKJ1-1 | GFTFSGSA | QSVRSSY | IVVGSGNT | GAS | AAPYCSSTSCRDGFDI | QQYGR*****SPWT |

VDJ alignment was determined by IgBLAST. The CDR sequences of 58G6 and 13G9 were aligned. Distinct amino acids were marked in red. The 94^th^ amino acid on the CDRH3 of mAb interacting with S^E484^ is labelled with asterisk.
